# GENATATORs: ab initio Gene Annotation With DNA Language Models

**DOI:** 10.64898/2026.06.17.732686

**Authors:** Aleksei Shmelev, Artem Shadskiy, Yuri Kuratov, Mikhail Burtsev, Olga Kardymon, Veniamin Fishman

## Abstract

Inference of gene structure and location from genome sequences - known as *de novo* gene annotation - is a fundamental task in biological research. However, sequence grammar encoding gene structure is complex and poorly understood, often requiring costly transcriptomic data for accurate gene annotation. In this work, we benchmark current solutions and develop new methods of gene annotation. We show that pre-trained DNA language model (DNA LM) embeddings do not capture the features necessary for precise gene segmentation, and that task-specific fine-tuning remains essential. We comprehensively evaluate the impact of model architecture, training strategy, receptive field size, dataset composition, and data augmentations on gene segmentation performance. We revisit standard evaluation protocols, showing that commonly used per-token and per-sequence metrics fail to capture the challenges of real-world gene annotation. We introduce and theoretically justify new biologically grounded metrics, along with benchmarking datasets that better capture annotation quality. We show that fine-tuned DNA LMs outperform existing annotation tools, generalizing across species separated by hundreds of millions of years from those seen during training, and providing segmentation of previously intractable non-coding transcripts and untranslated regions of protein-coding genes. Our results thus provide a foundation for new biological applications centered on accurate gene annotation.

## 1 Introduction

The rapid development of DNA sequencing technologies, such as third-generation sequencing and Hi-C, has led to an exponential growth in the availability of genome assemblies across the tree of life. This genomic data is invaluable for fundamental research, biotechnology, and biomedicine, but raw DNA sequences alone are insufficient for most applications. To interpret these data, genomes must be annotated, enabling the identification of functional elements. Gene annotation is the most important here, since it identifies genes and reveals their structural elements, which are critical for almost all downstream applications.

A gene is a continuous subsequence of genomic DNA that serves as the template for transcription, the process by which RNA molecules are synthesized from DNA. Genes are directional, and their direction is defined collinear with the direction of RNA synthesis. Therefore, genes can appear in forward or reverse orientation relative to the reference genome (Figure 1A). In the genomes, approximately half of the annotated genes are in the forward orientation and half in the reverse.

**Fig. 1.**
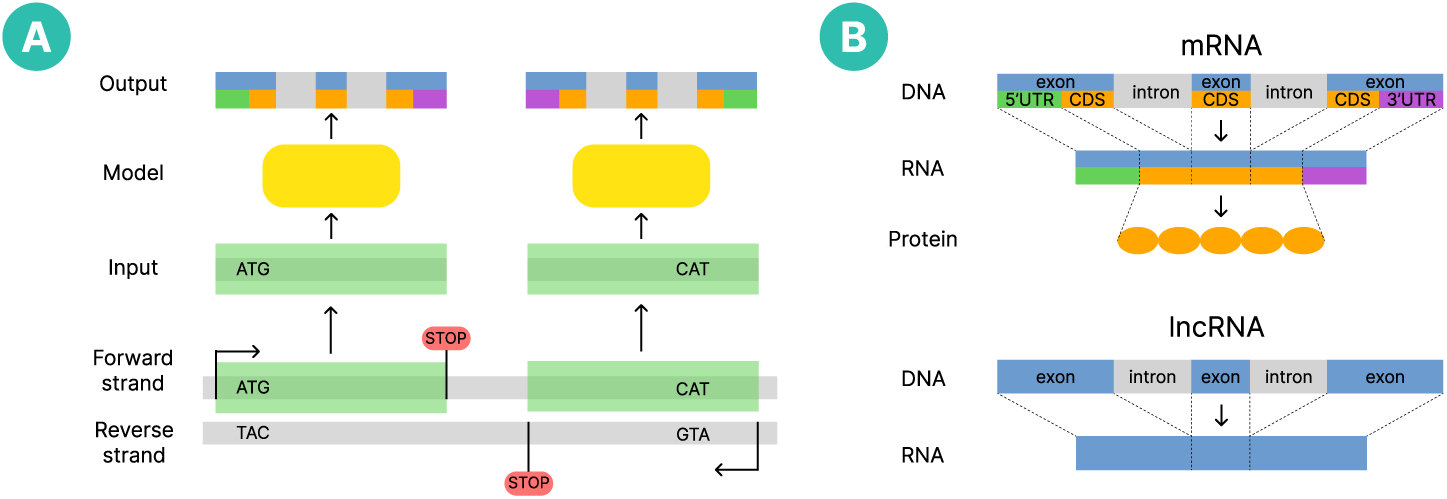
Gene structure and segmentation problem. Panel A shows transcript types in the dataset, where the model predicts all five classes but only intron and exon labels are relevant for lncRNAs, while all five are meaningful for mRNAs. Panel B illustrates that the model always receives DNA sequence from the forward strand (light green box) during training, yet these sequences may correspond to genes located on either strand.

The two largest gene classes are messenger RNAs (mRNAs) and long non-coding RNAs (lncRNAs). In the human genome, approximately 40.5% of genes are annotated as mRNAs and 35.2 % as lncRNAs, which is why this paper focuses only on these two classes. mRNAs encode proteins, and their sequence is segmented into exons and introns, with exons containing coding sequence (CDS) and untranslated regions (UTRs) at the 5^′^ and 3^′^ ends (Figure 1B). Translation of the CDS provides the amino acid sequence of proteins, each amino acid encoded by three CDS letters (codon). Thus, even a single nucleotide shift in an exon boundary can change all downstream codons. By contrast, lncRNAs lack CDS and do not produce proteins, but instead regulate diverse biological processes, including chromatin remodeling, immune response, viral defense, and cancer progression (16; 21). Untranslated regions of mRNAs are biologically important as well. Although they are not translated into proteins, UTRs influence transcript stability, translation efficiency, and localization (5). They may encode short functional peptides, and mutations in UTRs can be linked to human diseases (8). Thus, a complete view of gene structure requires accurate recovery not only of coding exons (CDS parts) but also of UTRs and non-coding genes.

Annotating lncRNAs and UTRs is a qualitatively different task compared to annotating mRNA CDS. Protein-coding gene fragments can often be recognized from conserved protein-coding fragments, while lncRNAs and mRNA gene UTRs lack such signals, evolve more rapidly, and are often expressed only in specific tissues, which makes their detection particularly challenging without additional evidence such as RNA-seq. Learning the sequence rules that govern transcription and protein synthesis should, in principle, enable the prediction of gene structure directly from DNA sequence. Methods that attempt this are known as *ab initio* gene predictors, yet in practice they underperform approaches that incorporate supplementary evidence beyond the genome sequence (19). Common sources of such evidence include gene annotations from closely related species and RNA-sequencing data from the target species (18). However, these resources are not consistently available across organisms or conditions, which sustains the demand for robust *ab initio* gene annotation methods that deliver high-quality results from sequence alone.

In this work, we address these gaps by applying DNA language models to gene segmentation and developing GENATATORs, a family of fine-tuned models specifically designed for *ab initio* annotation. Using biologically inspired metrics, justified by theoretical analysis and empirical validation, we demonstrate that pretrained DNA language model embeddings are insufficient for precise segmentation, making task-specific fine-tuning necessary. We then investigate how architecture, input context length, species diversity in training data, and augmentation strategies affect performance. Finally, we benchmark GENATATORs against existing methods and evaluate generalization on human and other species, showing that our models achieve state-of-the-art performance in gene segmentation due to the capacity to uncover previously untrackable lncRNAs and UTRs of mRNA, while maintaining comparable accuracy to the best existing tools on segmentation restricted to mRNA CDS.

## 2 Related Work

Early *ab initio* approaches relied on probabilistic models such as AUGUSTUS (22), which is based on HMMs that hardcode biological rules of gene grammar. These models capture statistical patterns of protein-coding genes, including the presence of a start codon to initiate CDS, a stop codon to terminate it, the absence of in-frame stops within the CDS, and canonical dinucleotides at splice junctions. Such models are effective for identifying protein-coding genes but fail to capture UTRs and lncRNAs (19). To address these gaps, deep learning methods have been introduced to learn gene segmentation rules from DNA sequences. Both Helixer and Tiberius combined CNNs with HMMs for gene annotation, with Tiberius achieving state-of-the-art accuracy on protein-coding gene annotation (11; 10). Although effective, these models remain constrained. Tiberius focuses on protein-coding genes and is not capable of retrieving UTRs or lncRNAs, and its CNN backbone is restricted to relatively short contexts (up to 10Kb) despite many human genes exceeding 30 Kb and spanning over 1 Mb.

Large DNA LMs have emerged as versatile backbones for genomic predictions (20; 9; 6; 14; 4; 24). Based on transformer or SSM architectures, they can be pre-trained on large genomic datasets. DNA LMs have matched or surpassed classical approaches across tasks such as splice-site prediction, promoter identification, and polyA signal detection. SegmentNT (7), a fine-tuned Nucleotide Transformer DNA LM (6) with a U-Net head, is a nucleotide-resolution classifier that outputs probabilities for each gene element directly from DNA sequence. Authors of SegmentNT also introduced variants of this model pretrained on expression data — SegmentBorzoi and SegmentEnformer. NTv3 (2) further scales context length to the megabase range. However, as we demonstrate below, classification performance on individual gene elements does not reliably reflect the accuracy of full gene reconstruction. Consequently, the utility of these models for real-world biological applications remains unclear.

Recently, AlphaGenome has been introduced as a foundation model of the genome that predicts multiple modalities from sequence, including RNA-seq, chromatin accessibility, and splicing-related outputs (1). In the splicing domain, it performs nucleotide-level classification of donor and acceptor sites, prediction of splice-site usage, and quantitative splice-junction prediction. While not being a gene annotation system, such splicing predictions of the model are directly relevant to exon–intron boundary detection and therefore to transcript assembly. Alongside these methods, several benchmarks have been proposed to assess gene annotation-related tasks. GUE (24) includes splice-site prediction. However, it assigns a single label to 400 bp input sequences, which makes it biologically irrelevant: gene annotation requires single-nucleotide precision in the detection of boundaries between gene elements. BEND (15) instead operates at the nucleotide level, but it uses short input sequences, relies on metrics that are not biologically rigorous, and does not evaluate critical elements such as UTRs or lncRNA genes. A detailed comparison between benchmarks developed in this work, BEND, and GUE is provided in Appendix A.

Building on these observations, it is clear that systematic evaluations of modern DNA LMs for full gene segmentation are still missing. In particular, SSMs have not been comprehensively benchmarked, and among transformer-based models, only a single context-extension method (17) has been applied to process genes longer than the default receptive field. A unified benchmark is therefore needed to clarify how modern DNA LMs perform on gene segmentation, especially for lncRNAs and UTRs that remain inaccessible to most existing tools.

## 3 Formal definition of the problem and metrics

We formalize gene segmentation as a multiclass nucleotide level classification task. The objective is to learn a function *f* that maps an input representation 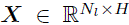 to an output label matrix 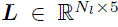, where *H* is the token embedding dimension, *N_l_* is the input length in nucleotides, and 5 is the number of target classes which are exon, intron, coding sequence (CDS), 5^′^ untranslated region (5-UTR), and 3^′^ untranslated region (3-UTR).

### 3.1 Segmentation scoring

#### Interval-level scoring

Segmentation performance can be assessed using conventional classification metrics such as precision, recall, f1-score, and PR-AUC computed per class at the nucleotide level. However, these metrics evaluate classification independently for each nucleotide and therefore may not capture biological dependencies between predictions. For instance, a misclassification of a single nucleotide within a megabase long gene has a negligible impact on the overall metric, while the same error can alter the interpretation of all down-stream sequence, since shifting a protein coding exon boundary by one nucleotide modifies all downstream trinucleotide blocks and yields a different amino acid sequence, a frame shift effect known in molecular biology.

To address this limitation, we use *interval level* segmentation scoring inspired by prior work (19). In this approach, a target interval is a continuous sequence of nucleotides with identical ground truth class labels. A predicted interval is counted as a true positive only when it has complete reciprocal overlap with a ground truth interval, which means that the predicted and true intervals coincide.

Formally, let the ground truth class label sequence be *L* = (*l*_1_*, l*_2_*, …, l_N__l_*). An interval *I_m_* = [*i, j*] is assigned to class *K* when *l_k_* = *K* for all *k* ∈ [*i, j*]. For each class *K*, 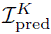 be the set of predicted intervals and let 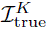 be the set of ground truth intervals. We compute the following quantities. True positives are the number of predicted intervals that exactly match a ground truth interval. False positives are the number of predicted intervals without an exact match 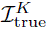. False negatives are the number of ground truth intervals that are not recovered in 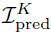

The final interval level f1-score for class *K* is

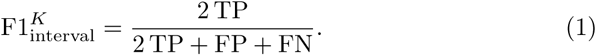

This metric penalizes biologically important segmentation errors and provides a realistic assessment of model performance.

#### Gene-level scoring

We also extend interval level scoring to evaluate the overall accuracy of gene structure prediction (defined as *gene level* metric). In gene level scoring, a gene is counted as a true positive only when all of its intervals are reconstructed correctly. Reference annotations may include multiple valid transcript structures for the same gene, known as transcription *isoforms*, which define different segmentations. To account for this ambiguity, we use a gene level rule that accepts a prediction as correct when the predicted interval set exactly matches the interval set of any annotated isoform of the target gene.

#### Gene-level scoring for CDS-only models

Many HMM-based models, including Tiberius and AUGUSTUS, rely on hard-coded parameters tailored to coding sequence identification, which makes them unable to detect exons that include untranslated regions. Therefore, for protein coding transcripts we report two gene level metrics, one where the complete exon structure is reconstructed (exon-mRNA) and one where only the coding sequence part is reconstructed (CDS-mRNA). For non-coding transcripts such as lncRNA, which have no CDS annotation, we compute gene level metrics using exon intervals only. To obtain an overall gene level score we sum the number of correctly predicted lncRNA genes by exon matching and the maximum of exon mRNA and CDS mRNA counts for protein coding genes, which allows a fair comparison across models

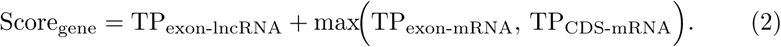

In Appendix B.1, we present a theoretical analysis that derives how the sensitivity of conventional PR-AUC and interval level metrics scales with boundary errors, justifying the need for the latter. This is followed by empirical evidence in Appendix B.2, where we demonstrate that relying on PR-AUC can lead to incorrect model rankings.

## 4 Experiments

*Input data* The training dataset consists of genes from all human chromosomes except 8, 20, and 21, which were held out for validation during training. When specified, we also included genes from all chromosomes of 39 additional mammalian species. All models were evaluated on human chromosome 20 from the T2T human assembly, since the human genome provides the most accurate annotation among all available species. This chromosome is part of the held-out set for several widely used models, including SegmentNT, Tiberius, and Helixer, enabling an unbiased comparison. For genes with multiple annotated isoforms, we selected a single isoform per gene with the longest cumulative length of exons. A detailed description of dataset preparation is provided in Appendix C.

*Models* We evaluated models representing different families of DNA LM architectures. From the SSM family, we included Evo2-1B (4) and Caduceus (with PH and PS modifications) (20). For Transformer-based models, we selected GENA-LM equipped with Recurrent Memory Transformer (RMT), capable of processing sequences comparable in length to complete genes (12). DNABERT-2, DNABERT-S, and similar architectures were not included due to the limited receptive fields. Additionally, we incorporated previously developed gene segmentation models based on the Nucleotide Transformer DNA LM: (SegmentNT and SegmentNT_multispecies), as well as models pretrained on gene expression data (SegmentEnformer and SegmentBorzoi), as well as classical models (HMM-based AUGUSTUS and the CNN&HMM hybrid Tiberius and Helixer), in the final benchmarks. Besides, we evaluated the recently released NTv3 and AlphaGenome models. However, we did not evaluate embeddings, reoptimize dataset preparation or training procedures for these models, as such studies have been reported previously (7; 10). We refer to the Appendix D Table A7 for the summary of all models benchmarked in this study.

For models operating at single-nucleotide resolution (Evo2 and Caduceus), we appended a linear projection layer of shape (*H,* 5) to map the model outputs to the five target classes. For non-single-nucleotide resolution models (Nucleotide Transformer, GENA-LM), token embeddings were upsampled by repeating each token representation to match its corresponding nucleotide span and further processed using a U-Net architecture as proposed in (7).

All models were trained using cross-entropy loss, and the best-performing checkpoint was selected based on exon-level f1-score on the validation set. Further details on model architectures and training protocols are provided in Appendix C.

### 4.1 Training on embeddings

DNA language models are expected to capture essential genomic features during pretraining. To evaluate whether gene-structure information can be extracted directly from frozen representations, we conducted experiments where the DNA LM weights were fixed and only a shallow classifier was trained. Specifically, we used a linear projection layer for models operating at nucleotide resolution (Evo2, Caduceus) and a U-Net decoder for the token-based GENA-LM (byte–pair–encoded inputs).

As shown in Appendix E, Table A8, none of the models produced embeddings containing sufficient information for accurate gene segmentation. The slightly higher performance of GENA-LM is likely attributable to the U-Net decoder, which, unlike the linear layer used in Evo2 and Caduceus, can aggregate local contextual signals.

To understand why pretrained models fail at segmentation, we analyzed final-layer hidden states on ten randomly selected human genes (six mRNA and four lncRNA) using Caduceus and GENA-LM. For GENA-LM, which uses BPE tokens, we expanded each token embedding uniformly across its nucleotide span to obtain one vector per base for both models. PCA projections of the Caduceus embeddings revealed four distinct clusters corresponding to nucleotide identity (A, C, G, T), rather than gene structure (Appendix F, Figure A2). GENA-LM embeddings formed diffuse clusters that also did not align with gene elements (Appendix F, Figure A1). This contrasts sharply with embeddings obtained after fine-tuning on the gene segmentation task described in Section 4.2, which show clear separation of gene elements (Appendix F, Figure A2). Quantitatively, fine-tuning increased the homogeneity of *k*-means (*k*=5) clusters with respect to exon, intron, CDS, 5^′^UTR, and 3^′^UTR labels from 0.003 to 0.583 for Caduceus and from 0.0 to 0.497 for GENA-LM.

These probing experiments should be interpreted in the context of our evaluation setup. In contrast to the BEND (15) benchmark, where authors employ task-specific trainable heads on top of frozen representations, our goal here is to assess what current pretrained encoders capture without relying on complex decoders. To keep the probing strictly aligned with this goal, we use only minimal heads (a linear layer for Evo2 and Caduceus, and a shallow U-Net for GENA-LM to enable nucleotide resolution), which reveals the information present in the embeddings themselves rather than what can be recovered by a powerful decoder.

Together, these results indicate that pretraining alone is insufficient to encode the features required for precise gene segmentation and that task-specific fine-tuning remains essential for achieving high segmentation accuracy.

### 4.2 Fine-tuning of DNA language models

We next conducted a series of fine-tuning experiments, where both the DNA LM parameters and the classification head were trainable. These experiments were designed to systematically investigate how model architecture and the biological information available during training influence gene segmentation performance.

As a baseline, we considered models trained on human genomic sequences with a model context length of 4,096 bp. Building on this setup, we explored the effect of extending the model context to 32 Kb, which provided a broader genomic window. We also examined whether expanding the training data to include genes from 39 additional mammalian species improved performance by leveraging evolutionary conservation, and we tested the impact of restricting the training set to protein-coding transcripts while excluding lncRNAs, so that the models were exposed only to sequences with well-defined coding structures. We additionally evaluated the effect of incorporating an auxiliary loss applied to intermediate model layers. Finally, in a complementary experiment, we evaluated training on multiple isoforms per gene vs using single representative isoform per gene in the baseline. In all experiments, we focused on Caduceus PS and Caduceus PH as representative SSMs, while GENA-LM served as the representative Transformer-based model, and we did not include Evo-1 and larger versions of Evo-2, since their size exceeded our available resources for running multiple fine-tuning experiments.

Our results (Table 1) indicate that increasing the input sequence length yields the most substantial improvement in segmentation performance, with approximately 1.6–2× gains across models. Incorporating multiple species into the training set improved performance by approximately 1.2–1.5×. Excluding lncRNAs from the training data resulted in improved CDS detection for both Caduceus models. However, this came at the expense of reduced lncRNA segmentation performance, although the decrease was not as pronounced. This observation suggests that the sequence grammar underlying non-coding transcripts can, to a large extent, be learned from protein-coding sequences. In contrast to CDS detection, we did not observe consistent improvements in exon segmentation for protein-coding genes when excluding lncRNA. Specifically, GENA-LM and Caduceus PS achieved a modest improvement of approximately 10%, whereas Caduceus PH exhibited a similar decrease in performance. Overall, we concluded that transcript filtering does not substantially improve training performance.

**Table 1.**
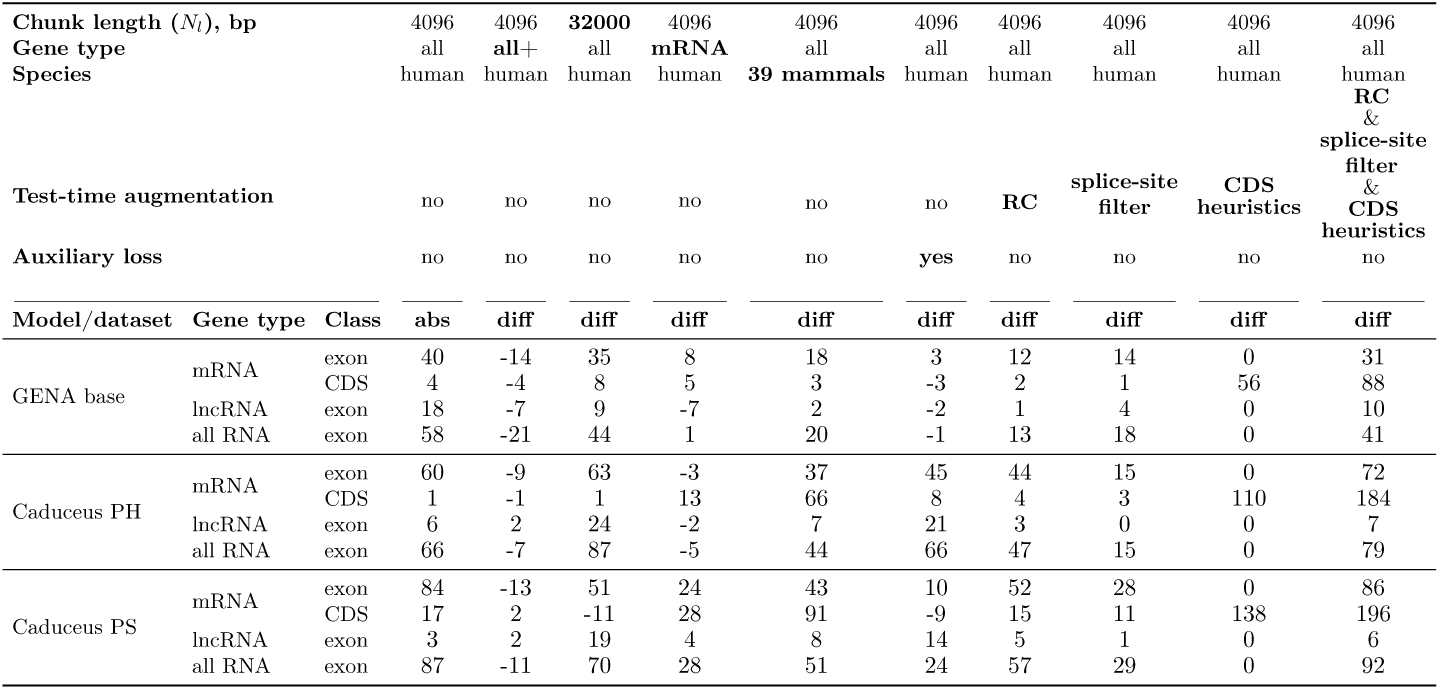
Gene-level performance metrics for dataset and model modifications: absolute number of correctly reconstructed genes (abs) and differences (diff) compared to baseline (presented in the first column). The gene type (all+) means that all isoforms of all genes were included in the dataset.

We also found that using multiple isoforms per gene slightly reduced accuracy, confirming that the single-isoform strategy remains preferable

Motivated by the auxiliary classifiers introduced in the Inception vision model, which use intermediate losses to improve optimization (23), we in particular evaluated the use of an auxiliary loss from the middle of the model. Concretely, for Caduceus we attached an independent linear projection head to the embeddings from layer 8 (Caduceus model has 16 layers), whereas for GENA-LM we attached an independent U-Net decoder to layer 6 (GENA-LM base total depth is 12 layers). During training, the losses from the intermediate and final heads were averaged to obtain the final training objective. As shown in Table 1, this modification substantially improved gene-level performance for Caduceus, with the strongest gains for Caduceus PH and consistent improvements for Caduceus PS, but it did not improve GENA-LM overall. This difference is likely related to the segmented nature of the Recurrent Memory Transformer architecture (12). In RMT the model has access only to the current segment and the memory carried over from preceding segments, but not to future tokens. As a result, auxiliary losses applied to earlier layers are less effective because those layers are required to make predictions before the model has traversed the full input sequence and accumulated the necessary long-range context. Based on this result, we used auxiliary loss in all final Caduceus experiments, but not in GENA-LM.

To investigate the biological features underlying model errors, we analyzed the precision and recall of exon detection, stratifying exon-intron boundary errors based on their flanking dinucleotide sequences (Appendix G, Figure A3). Although the frequency of predicted boundaries at each dinucleotide generally reflects the true distribution, we identified samples where dinucleotides flanking predicted boundaries never occur at boundary positions in the actual data. Explicitly excluding exons flanked by these “illegal” dinucleotides, designated as a “splice site filter” improves the performance of all models (Table 1).

As noted in the Introduction (Figure 1A), genes occur in both orientations relative to the reference genome, and for this reason, we apply a test-time reverse-complement (RC) augmentation in which each sequence is processed in its reference and RC orientations using average of the predictions as final output. As shown in Table 1, this approach yields substantial improvements in performance for all models. Notably, Caduceus PS, whose architecture explicitly enforces RC equivariance in the DNA input representation, still benefits significantly from test-time RC augmentation and achieves a ≈1.5× improvement in performance. This effect arises because sequences are segmented into fixed-size chunks and opposite orientations induce different chunkings, so averaging behaves like an ensembling. Furthermore, RC augmentation provides greater performance gains than applying a splice-site filter for both Caduceus models. To the best of our knowledge, this is the first study applying reverse-complement augmentation in the context of the gene segmentation task.

We additionally applied a CDS heuristic as a post-processing step for protein-coding transcripts. It concatenates the predicted exons, translates them in all reading frames, selects the longest resulting protein consistent with valid start and stop codons, and maps it back onto the predicted exons (Appendix I, Figure A4). As shown in Table 1, this substantially improves CDS reconstruction for mRNA transcripts, while not affecting lncRNA predictions.

Finally, we compared performance across model architectures. Consistent with previous benchmarks (20), Caduceus PS outperformed Caduceus PH in all experimental settings. The Transformer-based GENA-LM exhibited superior performance in lncRNA detection, whereas the SSM Caduceus detected a substantially higher number of protein-coding genes and achieved markedly better CDS segmentation compared to GENA-LM. We hypothesized that nucleotide counting is required to identify triplet-organized CDS. Whereas GENA-LM utilizes variable-length BPE tokens, making the counting task challenging, Caduceus employs single-nucleotide tokenization, which may explain improved performance for the CDS class. In contrast, GENA-LM consistently outperformed Caduceus in lncRNA segmentation, a task that is more challenging than mRNA for both models, and this advantage aligns with model capacity, since GENA base has approximately 120M parameters compared to 16M in Caduceus. When we trained the same base setup but with the larger 360M parameter GENA-LM, lncRNA segmentation performance improved by 25%, further highlighting the benefits of model scaling for this task (Appendix J, Table A12).

### 4.3 Scaling

To further improve model performance, we scaled and combined the features identified as most impactful for gene segmentation. Specifically, we increased the input sequence length to 250 Kb, utilized data from 39 mammalian species, and included all gene types in the training set. For the Transformer-based architecture, we employed a larger instance of GENA-LM with an increased number of parameters (GENA large), while for the SSM we used the Caduceus PS variant, which consistently demonstrated performance superior to Caduceus PH in our benchmarks.

At test time, we applied splice-site filtering, reverse-complement (RC) augmentation, and the CDS heuristic described earlier. We refer to the resulting models as GENATATORs, a DNA language model-based family of gene annotators.

After scaling, the Caduceus PS model achieved the best overall gene-level performance (Figure 2), outperforming GENA large and all other evaluated methods. In this final configuration, Caduceus PS was trained with auxiliary loss, longer context, and multispecies data, and this combination yielded best overall result. Contrary to our earlier observations in smaller-scale experiments, the scaled Caduceus PS model no longer shows a trade-off relative to GENA large, but instead achieves higher scores across all evaluated transcript categories, including CDS mRNA, exon mRNA, and lncRNA. GENA large remains the second-best model overall, whereas Tiberius retains competitive performance only on CDS-level protein-coding reconstruction and falls far behind on the full gene-level benchmark, which also evaluates exon-level mRNA reconstruction, including UTRs and lncRNA annotation.

**Fig. 2.**
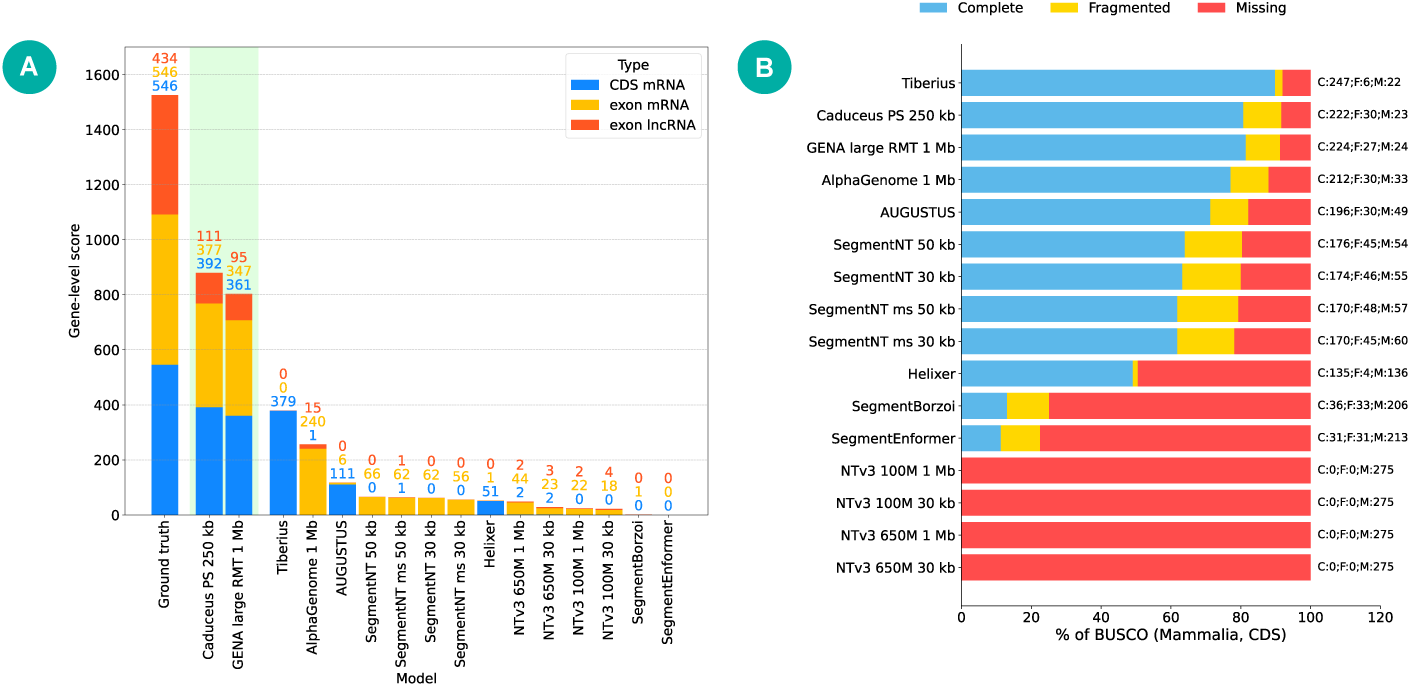
Comparison of gene segmentation performance across different annotation models. A. Gene-level metrics, GENATATORs are highlighted in green. Exon mRNA type also ensures correct UTRs prediction compared to CDS mRNA type. B. Mammalian BUSCO metrics.

### 4.4 Benchmarking GENATATOR against other gene-annotation tools

We evaluated the performance of the GENATATOR models in comparison with several state-of-the-art gene annotation tools, including the HMM-based AUGUSTUS (22), the CNN+HMM model Tiberius (10) and Helixer (11), the DNA LM-based SegmentNT (with variants trained on human-only and multispecies data) (7), and transformer-based models pretrained on gene expression, namely SegmentEnformer and SegmentBorzoi (7). We also included the recently developed AlphaGenome (1) and NTv3 (2).

We first compared models using the conventional PR-AUC metric (Appendix J, Table A13). According to this evaluation, GENATATORs outperform SegmentNT, SegmentBorzoi, and SegmentEnformer, with an improvement of about 10% between the best-performing GENATATOR and the best-performing SegmentNT.

We then assessed performance using the gene-level metrics described above, reporting results as the total number of correctly segmented genes across all evaluated categories, namely CDS mRNA, exon mRNA including UTRs, and lncRNA (Figure 2, with detailed metrics and model usage provided in Appendix I). Under this scoring scheme, GENATATORs identify substantially more genes than previously developed alternatives. In total, across these categories, GENATATORs show more than a threefold increase in the number of correctly segmented genes compared to Tiberius and AlphaGenome, and the gap to other models is even larger. Visual inspection of predicted gene structures reveals that lower-performing models such as SegmentNT frequently extend exon boundaries by several nucleotides, which in the case of mRNA leads to reading-frame shifts and consequently to biologically invalid truncated peptides. This observation underscores the importance of gene-level evaluation metrics for capturing biologically meaningful segmentation accuracy.

We attribute the improved performance of GENATATORs to a combination of training optimizations, including the use of multispecies data, extended input context lengths, and data augmentation strategies. As shown in Table 1, a basic training configuration with human-only data, a 4,096 bp input length, and no augmentations, or isolated modifications of this setup, produces results that are comparable to or worse than those achieved by SegmentNT.

GENATATORs also significantly outperform AUGUSTUS in the total number of correctly segmented genes. Stratifying performance by transcript type reveals that the Caduceus-based GENATATOR outperforms all other evaluated models, establishing a new state-of-the-art result in the total number of correctly segmented genes. In contrast, Tiberius outperforms the GENA-based GENATATOR in the number of correctly segmented protein-coding genes. However, Tiberius was developed primarily for protein-coding gene annotation and does not support broad annotation of lncRNA genes and UTRs within mRNA genes. This narrower annotation scope limits its overall gene annotation performance and leads to a lower total number of correctly segmented genes compared to GENATATOR.

The common metric for assessing the completeness of genome annotation is BUSCO (13). To compute BUSCO, the predicted exon-intron structure of a gene is used to generate an amino acid sequence, which is then compared to a set of proteins that are specific to a particular taxonomy group. The results of BUSCO are presented as a number of proteins that were identified from a selected dataset. These proteins are divided into two categories: Complete and Fragmented, where fragmented proteins have some segments missing.

Using the mammaliaspecific BUSCO dataset, GENATATORs recovered a comparable number of BUSCO orthologs to Tiberius, but with a larger fraction of fragmented genes. Caduceus identified 252 orthologs (222 complete and 30 fragmented), while GENA identified 251 orthologs (224 complete and 27 fragmented). In comparison, Tiberius identified 253 orthologs, including the highest number of complete BUSCOs (247) and only 6 fragmented genes. The three leading models, GENATATORs and Tiberius, were followed by AlphaGenome, which recovered 212 complete and 30 fragmented genes.

These results are consistent with the design of the BUSCO benchmark. BUSCO evaluates only a conserved subset of protein-coding transcripts. On chromosome 20, the RefSeq annotation contains 3,146 mRNA transcripts, whereas only 1,310 are represented in BUSCO. Accordingly, the advantage of GENATATORs is more evident in our transcript-level reconstruction metrics than in BUSCO, while Tiberius appears biased toward more conserved isoforms.

Other models, including SegmentNT, SegmentBorzoi, SegmentEnformer and NTv3 showed substantially lower BUSCO recovery rates, consistent with their lower gene level segmentation performance. These results further reinforce the conclusion that conventional classification metrics such as PR-AUC are poor proxies for evaluating the biological utility of the models.

We next investigated whether segmentation errors made by different tools are shared or model-specific. Shared errors would suggest the presence of genes with structural features that are out-of-distribution relative to the training data, while model-specific errors would indicate that each tool fails on a unique subset of genes. To explore this, we analyzed the overlap of correctly segmented genes among the three top-performing models: the two GENATATOR variants and Tiberius. As shown in Figure 3, there is a substantial intersection of correctly segmented genes across all models, supporting the hypothesis that certain genes present a challenge to all tools. At the same time, each model also segments a distinct subset of genes not correctly annotated by the others. In comparisons between GENATATORs and Tiberius, the unique gene set recovered by GENATATORs is largely composed of lncRNAs, which Tiberius is not designed to annotate. Meanwhile, the unique gene set recovered by Tiberius is likely enriched in conserved genes, which may explain the higher number of complete BUSCO genes recovered by this model (Figure 2B). These findings suggest that model ensembling is currently the most effective strategy for maximizing gene annotation coverage across both coding and non-coding transcripts.

**Fig. 3.**
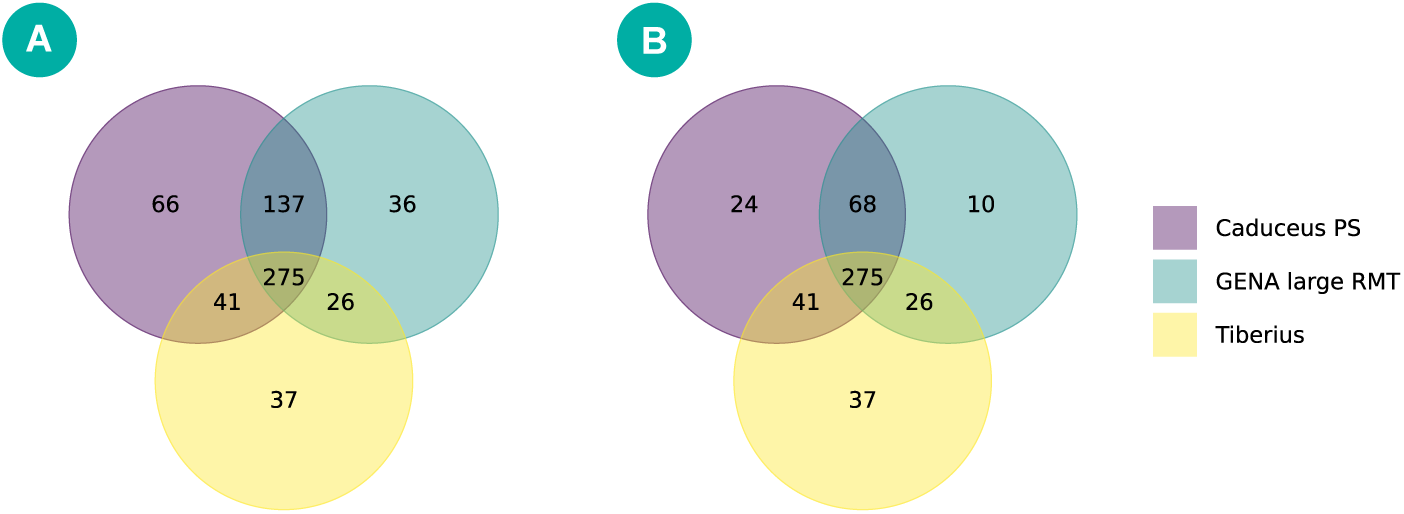
Each model provides unique set of annotated genes, yet large portion of errors are shared accross models. Overlap of correctly segmented genes shown for protein-coding and lncRNA genes together (A), and for protein-coding genes only (B).

Overall, our results position GENATATORs as state-of-the-art models for gene annotation, with particular strength in the detection of non-coding genes and UTRs. The computational resource requirements of our models are described in Appendix H.

### 4.5 GENATATORs generalize across unseen species at large evolutionary distances

A key application of *ab initio* gene predictors is the annotation of genomes from previously unannotated species. To evaluate the cross-species generalization of our models, we first evaluated performance using gene-level metric on two evolutionarily remote species representing different kingdoms of life: the flowering plant *Arabidopsis thaliana* (GCF_000001735.4) and the budding yeast *Saccharomyces cerevisiae* (GCF_000146045.2) (Table 2). At the nucleotide level, there is effectively no sequence homology between their genes and those of mammalian species included in the training dataset, and thus the models had never encountered any comparable sequences during training. Despite this extreme divergence, the models retained reasonable accuracy. For *A. thaliana*, both Caduceus PS and GENA large showed strong cross-species generalization, with Caduceus PS achieving the best exon-level reconstruction overall and substantially outperforming Tiberius, including its specialized version, as well as AUGUSTUS. In CDS reconstruction for *A. thaliana*, however, AUGUSTUS achieved the best result, whereas both versions of Tiberius remained well below Caduceus PS and GENA large. For *S. cerevisiae*, whose compact genome lacks spliceosomal introns, Caduceus PS again achieved the best performance and outperformed all baselines, whereas Tiberius and its specialized version failed to recover any genes. NA entries in the lncRNA row of Table 2 indicate the absence of annotated lncRNAs in the reference genome.

**Table 2.**
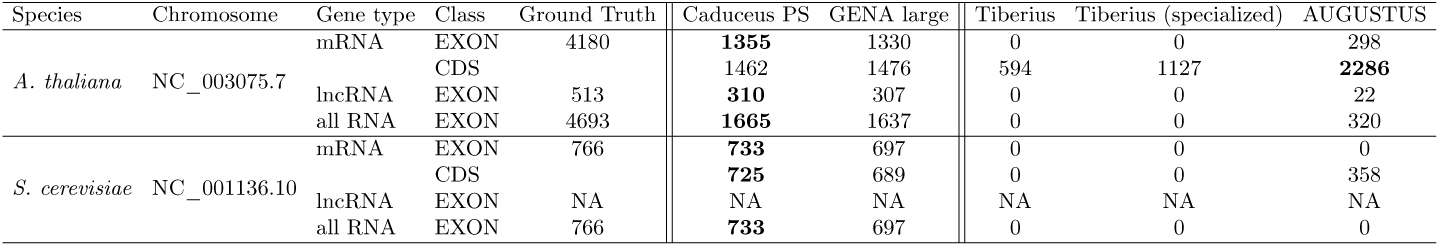
Gene-level performance of different models on evolutionarily distant species. Tiberius (specialized) column denotes the use of model weights pretrained on the phylogenetic group corresponding to the target organism (Eudicotyledons for *A. thalian* and Fungi for *S. cerevisiae*).

| Species | Chromosome | Gene type | Class | Ground Truth | Caduceus PS | GENA large | Tiberius | Tiberius (specialized) | AUGUSTUS |
| --- | --- | --- | --- | --- | --- | --- | --- | --- | --- |
| *A. thaliana* | NC_003075.7 | mRNA | EXON | 4180 | **1355** | 1330 | 0 | 0 | 298 |
|  |  |  | CDS |  | 1462 | 1476 | 594 | 1127 | **2286** |
|  |  | lncRNA | EXON | 513 | **310** | 307 | 0 | 0 | 22 |
|  |  | all RNA | EXON | 4693 | **1665** | 1637 | 0 | 0 | 320 |
| *S. cerevisiae* | NC_001136.10 | mRNA | EXON | 766 | **733** | 697 | 0 | 0 | 0 |
|  |  |  | CDS |  | **725** | 689 | 0 | 0 | 358 |
|  |  | lncRNA | EXON | NA | NA | NA | NA | NA | NA |
|  |  | all RNA | EXON | 766 | **733** | 697 | 0 | 0 | 0 |

Same results were obtained when we excluded all genes with detectable protein-level similarity to mammals, to ensure that models can not find homology even after internally translating DNA to amino acid code. Under this stringent setting, GENATATORs reconstructed more than twice as many genes as AUGUSTUS, despite the latter being run with a species-specific profile (Appendix K). Thus, although not tuned for plants or fungi, the models were able to produce useful first-pass annotations in such genomes, providing strong evidence that their capabilities extend beyond mere memorization of homologous patterns.

In addition to this extreme test, we benchmarked the models across a spectrum of animal species, ranging from primates closely related to humans to distant lineages such as insects (Figure 4). The ranking of methods remained largely consistent across species. DNA language models outperformed classical annotation tools, with Caduceus PS achieving the best overall performance and GENA large usually ranking second. In contrast, Tiberius and AUGUSTUS performed substantially worse on evolutionarily distant organisms. For protein-coding genes, segmentation accuracy decreased with evolutionary distance from human, whereas lncRNA detection was lower in mammals but higher in several distant taxa. In most species, Caduceus PS matched or exceeded GENA large.

**Fig. 4.**
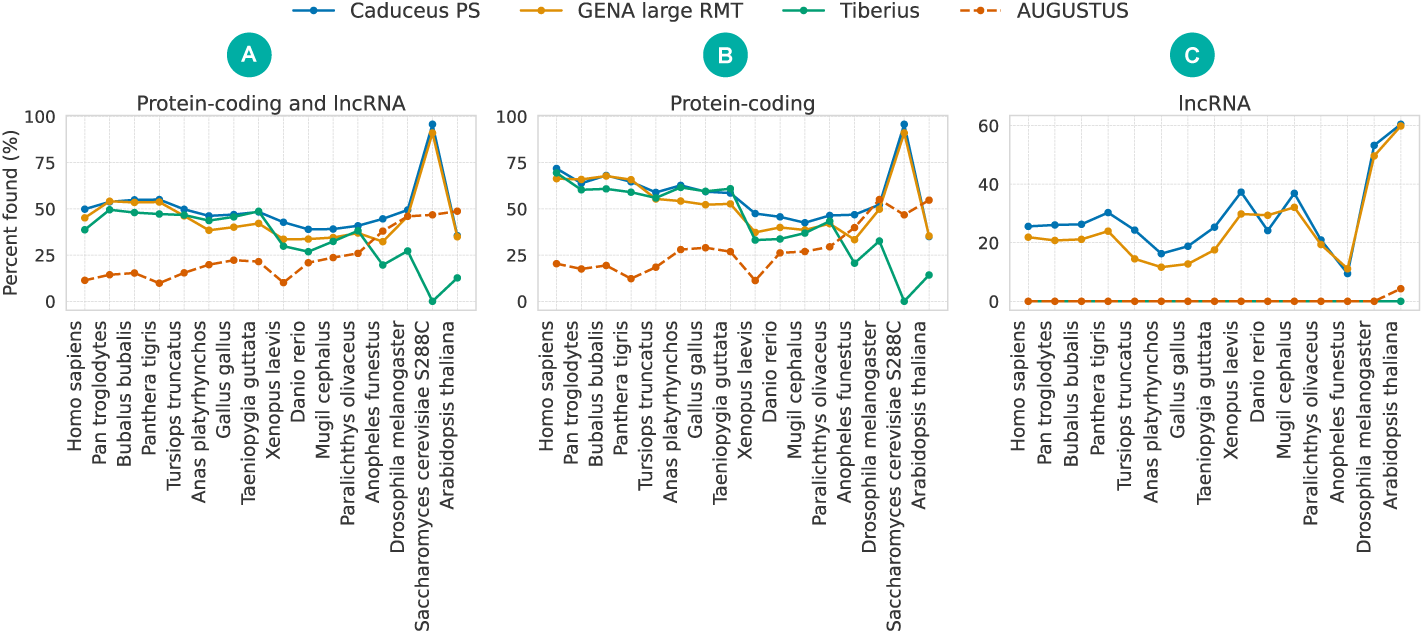
GENATATORs generalize to previously unseen species. Performance of the models in human and 13 other species for all (A), protein-coding (B), and lncRNA (C) genes. *S. cerevisiae* was excluded from panel (C) because it lacks lncRNA genes. See Appendix J Tables A16 for more information on metrics.

## Conclusions

In this work, we comprehensively evaluated the utility of DNA LMs for the gene segmentation task. We show, both theoretically and empirically, that interval level metrics better reflect biological relevance than conventional token level classifiers and introduce dedicated benchmark to score gene segmentation models. We demonstrated that embeddings from pretrained DNA LMs do not contain sufficient information for accurate gene segmentation. However, by identifying optimal training regimes, datasets, augmentations, and output filters, we enabled efficient fine-tuning and inference of gene structure. We further showed that scaling DNA LMs under these conditions substantially improves performance, leading to state-of-the-art results.

We found that sensitivity to different functional gene elements, such as CDS and UTRs, varies across DNA LM architectures. Nonetheless, all evaluated DNA LMs were capable of detecting lncRNA genes, which remain inaccessible to current state-of-the-art tools such as Tiberius.

Furthermore, GENATATORs, our fine-tuned DNA LM-based models, generalize effectively to unseen species across large evolutionary distances. These results highlight the potential of DNA LMs to serve as powerful tools for *de novo* genome annotation in a wide range of biological and evolutionary studies.

## Acknowledgments

Support from the Basic Research Program of HSE University is gratefully acknowledged (HSE-BR-2025-020).

## Code availability

HuggingFace versions of the datasets and models, in PyTorch, can be found via HuggingFace at https://huggingface.co/collections/ AIRI-Institute/genatator. The gene-segmentation benchmark is available via HuggingFace at https://huggingface.co/spaces/AIRI-Institute/genatator-ab-initio-segmentation-leaderboard.

## Limitations

While GENATATORs demonstrate strong performance in benchmarking studies, their accuracy remains far from perfect. Currently, only approximately 30–40% of all human genes can be correctly segmented by any of the models evaluated in this study.

Another limitation lies in gene discovery. Although gene segmentation is a critical component of genome annotation, complete annotation also requires accurate identification of gene boundaries, including non-coding untranslated regions (UTRs), which remains challenging for all evaluated tools.

Finally, the poor results observed in our embedding-only training experiments highlight a fundamental limitation of current DNA language models: they do not capture gene structure during the pretraining phase. This underscores the need for architectural or training paradigm improvements in future DNA LM development.

