## Supplementary Material for "GENATATORs: ab initio Gene Annotation With DNA Language Models"

#### Appendix A. Differences between our benchmark and others.

This appendix compares our benchmark with BEND (15) and with GUE introduced alongside DNABERT-2 (24), focusing on input length coverage, task granularity, and the biological meaning of reported metrics. Table A1 summarizes the design choices in each suite, and Table A2 reports human training-set lengths that illustrate coverage differences.

**Table A1.** Design comparison of benchmarks.

| **Benchmark** | **Input scope** | **Typical length** | **Granularity** | **Evaluation scope** | **Metrics** |
|---|---|---|---|---|---|
| GUE (24) | short sequences | 70–1000 bp; splice sites 400 bp; GUE+ 5–10 kb | sequence-level | local classification tasks | task-specific (MCC / F1) |
| BEND (15) | gene snippets | up to 13 kb | nucleotide-level | nucleotide classification of gene-structure labels; no full-gene segmentation; no UTR / lncRNA | MCC only |
| Ours | full genes via tiling | train 4 096 or 32 k or 250 k nt; full gene length evaluation | nucleotide-level | end-to-end segmentation with full gene reconstruction; with UTR and lncRNA | interval-, gene-level |

A key difference is context length. As summarized in Table A2, our training data span substantially longer transcripts than BEND, preserving the long tail of gene length distribution. In fact, 17,737 human transcripts in our set exceed 13,000 nt, whereas BEND truncates at this length. In addition, sequence-level suites such as GUE emphasize short-range classification and report scores that do not capture boundary accuracy, while BEND, although nucleotide-level, uses metrics that are not biologically rigorous for full gene structures and does not assess UTRs or lncRNA genes. By contrast, our evaluation targets complete gene structures with interval- and gene-level metrics. A detailed analysis of metric sensitivity appears in Appendix B.

*Benchmarking on BEND* For comparability, we also report results on BEND. Unlike the probing setup in the original BEND paper that assesses the quality of the embeddings in different pretrained models, we fine-tuned our models until convergence using the official train, validation, and test splits. This decision was deliberate: BEND compared all models against AUGUSTUS, which is a trained HMM genome annotation tool (it saw all human genes in the BEND benchmark during training). To ensure fairness we therefore also trained our models. Because sequences in BEND are short, all of our models can handle the full length of

**Table A2.** Statistics of training datasets for BEND and our benchmark (human).

| Dataset | Transcripts | Mean (nt) | Median (nt) | 95th perc. (nt) | Max (nt) |
|---|---|---|---|---|---|
| BEND (human) | 4,783 | 7,474 | 7,355 | 12,414 | 13,000 |
| Ours (human) | 33,367 | 37,366 | 14,651 | 176,543 | 250,000 |

each sample, so no chunking was applied at either training or validation. The reported metric is MCC, as specified in the BEND paper.

**Table A3.** BEND gene-finding results (MCC) with fine-tuned models using official splits and full-sequence inference.

| Model | MCC |
|---|---|
| Caduceus PS | **0.83** |
| AUGUSTUS | 0.80 |
| Caduceus PH | 0.72 |
| GENA base | 0.65 |

*Comparison of pur benchmark with G3PO and Tiberius approaches* The G3PO benchmark (19) is constructed from 1,793 UniProt proteins grouped into twenty orthologous families selected to represent complex protein-coding genes across 147 species. For each protein, the corresponding genomic locus and exon map are retrieved from Ensembl, and evaluation is carried out at nucleotide, exon, and protein levels against a single reference protein per gene. Consequently, G3PO covers only protein-coding genes, excludes lncRNA, and does not assess complete gene structure across multiple transcript isoforms.

`Tiberius` (10) is trained on mammalian protein-coding genes and uses convolutional and recurrent layers combined with a differentiable HMM. To obtain unambiguous labels, only the transcript with the longest coding sequence is retained for each gene, and evaluation is performed against this single coding isoform. As a result, exon- and gene-level metrics for Tiberius are computed relative to one reference isoform rather than across the full isoform set.

In contrast, our benchmark evaluates complete exon structures for all supported transcript types, including UTR exons, coding exons, and exons of lncRNAs. A prediction is counted as correct only when the full set of predicted exons matches the exon set of at least one annotated isoform, which allows transcripts containing both coding and non-coding segments to be evaluated faithfully. Together with CDS-based metrics comparable to those used in (19), our interval- and gene-level metrics provide a more biologically aligned assessment of complete gene reconstruction for both coding and non-coding genes.

### Appendix B. PR-AUC sensitivity and supporting evidence.

#### B.1 Theoretical evidence

Per nucleotide metrics such as precision, recall, f1 and PR-AUC treat each base independently, which can hide small local mistakes that have large biological impact. We provide theoretical evidence of this discrepancy between nucleotide and interval level metrics using a binary setup with two mutually exclusive classes, exon coded as 1 and intron coded as 0. For a single gene containing $p$ positive exon bases and $n$ negative intron bases with positive scores $s_i$, PR-AUC equals Average Precision and can be written using the ranks of positives in the list sorted by $s_i$ in descending order

$$\text{PR-AUC} = \text{AP} = \frac{1}{p} \sum_{k \in R_+} \Pr(k), \qquad \Pr(k) = \frac{\#\text{positives in top } k}{k}, \tag{3}$$

where $R_+$ is the set of positions in the sorted list that are occupied by positives. This depends only on the ordering of scores, so any monotone transformation that preserves order keeps PR-AUC unchanged.

We now carry one simple example through the derivation so that each step is explicit. Consider a short gene with a single exon block followed by an intron block. The targets and baseline scores are

$$y = \big[1,\ 1,\ 1,\ 1,\ 0,\ 0,\ 0,\ 0\big] \quad \text{and} \quad s = \big[0.99,\ 0.95,\ 0.92,\ 0.91,\ 0.40,\ 0.35,\ 0.31,\ 0.20\big].$$

This is a good prediction because exons receive higher scores than introns. The scores are already in descending order, so the cumulative number of exons in the top $k$ positions is

$$T(1) = 1,\ T(2) = 2,\ T(3) = 3,\ T(4) = 4,\ T(5) = 4,\ T(6) = 4,\ T(7) = 4,\ T(8) = 4,$$

and the corresponding precision values are

$$\Pr(1) = \tfrac{1}{1},\ \Pr(2) = \tfrac{2}{2},\ \Pr(3) = \tfrac{3}{3},\ \Pr(4) = \tfrac{4}{4},\ \Pr(5) = \tfrac{4}{5},\ \Pr(6) = \tfrac{4}{6},\ \Pr(7) = \tfrac{4}{7},\ \Pr(8) = \tfrac{4}{8}$$

Average Precision averages these precision values only at the positive positions $k \in \{1, 2, 3, 4\}$, hence

$$\text{AP} = \frac{1}{4}\Big(1 + \frac{2}{2} + \frac{3}{3} + \frac{4}{4}\Big) = 1. \tag{4}$$

If we apply a monotone change to all scores, for example $s \mapsto s^2$ or $s \mapsto s + 5$, the order does not change and equation 4 remains the same, which illustrates the order invariance of PR-AUC in equation 3.

We now introduce a boundary error at the exon edge before sorting and we make the modification explicit. Keep the targets $y$ fixed and lower the scores of the last two exon bases so that they fall below all intron scores. Define the modified score vector

$$\tilde{s} = \big[0.99,\ 0.95,\ \underline{0.19},\ \underline{0.18},\ 0.40,\ 0.35,\ 0.31,\ 0.20\big],$$

where the underlined entries mark the two exon bases affected by the boundary error. This change is applied before sorting by score. After sorting $\tilde{s}$ in descending order, the new score order is

$$\tilde{s}_{\text{sorted}} = \big[0.99,\ 0.95,\ 0.40,\ 0.35,\ 0.31,\ 0.20,\ 0.19,\ 0.18\big],$$

and the corresponding sorted labels become

$$y'_{\text{sorted}} = \big[1,\ 1,\ 0,\ 0,\ 0,\ 0,\ 1,\ 1\big].$$

Thus the two undemoted exons stay at ranks 1 and 2, the four introns occupy ranks 3 through 6, and the two demoted exons move to ranks 7 and 8. The cumulative positives for the modified order are

$$T'(1) = 1,\ T'(2) = 2,\ T'(3) = 2,\ T'(4) = 2,\ T'(5) = 2,\ T'(6) = 2,\ T'(7) = 3,\ T'(8) = 4$$

and the Average Precision after the error averages the precision values at the positive ranks $1, 2, 7, 8$

$$\text{AP}' = \frac{1}{4}\Big(\frac{1}{1} + \frac{2}{2} + \frac{3}{7} + \frac{4}{8}\Big) = \frac{1}{4}\Big(1 + 1 + \frac{3}{7} + \frac{1}{2}\Big) = \frac{41}{56} \approx 0.7321.$$

We now connect this explicit computation with the general formula. In the general case with $p$ exon nucleotides and $n$ intron nucleotides, if $\delta$ exon bases near the boundary are lowered below all intron scores before sorting, the sorted list contains $p - \delta$ exons first, then $n$ introns, then the $\delta$ demoted exons. The $r$th demoted exon occupies rank

$$k_r \ = \ n + (p - \delta) + r \quad \text{for} \quad r = 1, \ldots, \delta,$$

because the top contains $p - \delta$ undemoted exons and $n$ introns before the first demoted exon appears. At rank $k_r$ the prefix contains $(p - \delta) + r$ exons, so its precision equals

$$\Pr(k_r) \ = \ \frac{p - \delta + r}{n + p - \delta + r}.$$

All remaining $p - \delta$ exons at ranks 1 through $p - \delta$ have precision 1. Plugging these two groups into equation 3 gives the exact PR-AUC after the boundary error

$$\text{PR-AUC}' = \frac{1}{p}\Big[(p - \delta) \cdot 1 + \sum_{r=1}^{\delta} \frac{p - \delta + r}{n + p - \delta + r}\Big]. \tag{5}$$

For the example with $p = 4$, $n = 4$ and $\delta = 2$ this yields

$$\text{PR-AUC}' = \frac{1}{4}\Big[2 \cdot 1 + \frac{3}{7} + \frac{4}{8}\Big] = \frac{41}{56},$$

which is exactly the value computed from the sorted example above.

The corresponding loss is

$$\begin{aligned}\Delta\text{PR-AUC} = 1 - \text{PR-AUC}' &= \frac{1}{p}\sum_{r=1}^{\delta}\Big(1 - \frac{p-\delta+r}{n+p-\delta+r}\Big)\\ &= \frac{1}{p}\sum_{r=1}^{\delta}\frac{n}{n+p-\delta+r}\\ &\le \frac{1}{p}\sum_{r=1}^{\delta}\frac{n}{n+1} = \frac{\delta\, n}{p\,(n+1)} \le \frac{\delta}{p}.\end{aligned} \tag{6}$$

The last two inequalities hold because each denominator satisfies $n+p-\delta+r \ge n+1$, hence each summand is at most $n/(n+1) < 1$, so the sum of $\delta$ such terms is at most $\delta\, n/(n+1) < \delta$, and dividing by $p$ yields the stated bound $\Delta\text{PR-AUC} \le \delta/p$.

Under the same error the interval and gene views behave differently. If the gene has $m$ true exon intervals and the boundary of one interval moves by one base, that interval no longer matches exactly. True positives drop from $m$ to $m-1$ and at least one false positive and one false negative appear. Substituting into Eq. equation 1 yields

$$\text{F1}^{\text{exon}}_{\text{interval}} = \frac{2(m-1)}{2(m-1)+2} = 1 - \frac{1}{m}. \tag{7}$$

Define the interval drop as the difference between the perfect and the post error score. With one boundary shift that breaks exactly one interval and introduces exactly one false positive and one false negative, the drop is

$$\Delta\text{F1}^{\text{exon}}_{\text{interval}} = 1 - \Big(1 - \frac{1}{m}\Big) = \frac{1}{m}, \tag{8}$$

and it can be larger if the prediction creates additional spurious or missed intervals.

At gene level the same single boundary shift breaks the exact match for all isoforms, so the gene contributes 1 before the error and 0 after

$$\Delta\text{Score}_{\text{gene}} = 1. \tag{9}$$

Given equation 6, equation 8 and equation 9, the sensitivity fractions for the same local error satisfy

$$\frac{\Delta\text{PR-AUC}}{\Delta\text{Score}_{\text{gene}}} \le \frac{\delta}{p} \quad \text{and} \quad \frac{\Delta\text{PR-AUC}}{\Delta\text{F1}^{\text{exon}}_{\text{interval}}} \le \frac{\delta}{p}\, m. \tag{10}$$

With $m$ fixed and $p$ large the right-hand sides are small. Therefore, given the same boundary mistake, PR-AUC changes by at most $\delta$ over $p$ and becomes negligible on long exons, while the interval score and the gene score incur fixed drops per affected interval and per affected gene.

##### B.2 Empirical evidence

We complement the theory with experiments scoring models with PR-AUC and interval level metrics (Table A4). These results show that model ranking depends on the metrics used.

**Table A4.** Why gene level metrics matter, comparison of mean PR-AUC and fully reconstructed genes

| model | PR-AUC mean | gene level all |
|---|---|---|
| Caduceus PH 32 kb | 0.7728 | 276 |
| Caduceus PS 32 kb | 0.7834 | 267 |
| GENA-LM 1M | 0.7815 | 442 |

Both `Caduceus` variants exceed `GENA-LM` by PR-AUC mean, yet they reconstruct about three times fewer genes, since 276 versus 442. Across all models, the spread in mean PR-AUC is about 0.16, for example, `Caduceus PS` 0.8315, `SegmentNT` 0.7131, `SegmentEnformer` 0.5329 (Appendix J Table A13), while the difference in fully reconstructed genes ranges from 0 to 442. With these numbers in mind, optimizing only PR-AUC during early experiments can reward architectures that seem promising while failing to assemble biologically valid transcripts, which slows progress.

We further trained models on a human gene set with the same labels but one label per BPE token and varied input length from 4k BPE tokens which is approximately 32k nucleotides to 32k tokens, which is approximately 250k nucleotides.

**Table A5.** Effect of input length and output granularity on PR-AUC mean and gene level all for `GENA large`

| setting | input length nt | PR-AUC mean | gene level all |
|---|---|---|---|
| 4 k | $\approx$ 32k | 0.6787 | 16 |
| 16 k | $\approx$ 128k | 0.7304 | 22 |
| 32 k (BPE) | $\approx$ 250k | 0.7464 | 24 |
| 32 k (nucleotide, human) | $\approx$ 250k | 0.7459 | 109 |

Mean PR-AUC differs by about 0.0677 between the 4k and 32k BPE models, yet the gene level score rises from 16 to 24 which is a factor of about 1.5. Switching from BPE outputs to nucleotide outputs by stacking a U-Net on top of the trained model changes PR-AUC from 0.7464 to 0.7459, while the number of fully reconstructed genes increases by 85 which is a factor of about 4.5. With the arguments provided in Section B.1 and these empirical trends, we get that context length and boundary precision both matter for transcript assembly and that interval and gene level evaluation is needed when developing annotation models.

#### Appendix C. Dataset preparation, model training and architecture details

The dataset was constructed using the human genome assembly GCF_009914755.1. Chromosomes 8, 20 and 21 were designated as the validation set, but only chromosome 20 was used to compute final metrics for computational efficiency. We did not use a separate test set. The dataset contains all mRNA and lncRNA genes, and all sequences were exclusively from the forward strand. The dataset was filtered via selecting one representative transcript per gene, choosing the longest transcript available. Only transcripts with a length of up to 250 Kb were included.

Below we provide details of modifications in dataset, training regime or architecture for specific models:

1. For the mRNA-only dataset, we selected samples corresponding exclusively to protein-coding genes from the original dataset.
2. For the multispecies dataset, we processed data for 39 species (38 plus human) using the same strategy as for human samples. The list of species is provided in Table A6. It's important to note that only the human genome is fully assembled, therefore samples from other species containing 'N' characters (indicating unknown sequences) were excluded.
3. All models were trained using flash attention support (if supported by the model) to improve computational efficiency.
4. For training BPE-based `GENA` models at nucleotide-level resolution, embeddings derived from the token-level models were employed, omitting memory, CLS, and SEP tokens. The primary distinction between handling embeddings from `GENA-LM` versus other models arises from `GENA-LM`'s use of BPE tokens, necessitating additional steps before U-Net usage, whereas models like `Caduceus` and `Evo2` already operate directly at nucleotide resolution. Specifically, for `GENA-LM`, token embeddings were upsampled, meaning each embedding was replicated according to how many nucleotides it covered. Subsequently, nucleotide-specific embeddings (one per nucleotide type, a total of four different learnable embeddings) were concatenated to these upsampled token embeddings. For computational efficiency, those embeddings were segmented into non-overlapping chunks of 8192 base pairs (along sequence length axis), which were individually fed into the U-Net model. In contrast, for models that can directly utilize nucleotide resolution, we simply included an additional fully connected layer to convert embeddings into class probability vectors.
5. A learning rate of $5 \times 10^{-5}$ and weight decay of $1 \times 10^{-4}$ with AdamW optimizer was discovered to be the optimal trade-off between prediction accuracy (particularly for splice site boundary detection) and convergence speed, as lower values adversely impacted prediction quality.
6. Training of each model was performed on 8 Nvidia GPUs (either A100 or H100), except for Evo2, which specifically required Nvidia H100 GPUs due to compatibility constraints (GPU compatibility $> 8.9$). All models were

trained until convergence was observed using an exon-level validation metric. Typically, training with frozen embeddings required approximately half a day, while low-scale finetuning took about two days, with slight variations depending on the specific model. It took us one week to train the final models presented in our benchmark section.

7. In training and internal validation we do not always take nucleotides from the beginning of a gene. Instead, we choose a random starting position and extract at most $N$ nucleotides to the right, where $N$ is the model's context length (4096, 32k, or 250k as reported in the main text). We also ensure that the selected subsequence is at least 512 nucleotides long, so that the model always receives enough context. Each gene contributes a single subsequence of this form, with no splitting. Metrics computed in this setup, such as PR-AUC and interval level scores, are used only to select the best checkpoint for later evaluation.
8. For the final validation reported in the paper, we evaluate complete genes. Here, sequences are divided into non-overlapping chunks of the same length that the model was trained on. Predictions are made for each chunk, then concatenated to recover the full gene, and metrics are calculated on the full-gene predictions. This guarantees consistency with training while still allowing evaluation of arbitrarily long genes.

**Table A6.** List of genomic assemblies used to create the multispecies training dataset. List of genomic assemblies used to create the multispecies training dataset. Assembly names correspond to the annotation and genome names. The annotation files were obtained from NCBI.

| Assembly | Species |
|---|---|
| GCF_000952055.2 | Aotus nancymaae |
| GCF_002263795.3 | Bos taurus |
| GCF_000767855.1 | Camelus bactrianus |
| GCF_000002285.3 | Canis lupus familiaris |
| GCF_000151735.1 | Cavia porcellus |
| GCF_001604975.1 | Cebus imitator |
| GCF_000283155.1 | Ceratotherium simum simum |
| GCF_000276665.1 | Chinchilla lanigera |
| GCF_000260355.1 | Condylura cristata |
| GCF_002940915.1 | Desmodus rotundus |
| GCF_000151885.1 | Dipodomys ordii |
| GCF_002288905.1 | Enhydra lutris kenyon |
| GCF_000308155.1 | Eptesicus fuscus |
| GCF_000002305.2 | Equus caballus |
| GCF_018350175.1 | Felis catus |
| GCF_000247695.1 | Heterocephalus glaber |
| GCF_009914755.1 | Homo sapiens |
| GCF_000236235.1 | Ictidomys tridecemlineatus |
| GCF_000280705.1 | Jaculus jaculus |
| GCF_000001905.1 | Loxodonta africana |
| GCF_001458135.1 | Marmota marmota |
| GCF_000165445.2 | Microcebus murinus |
| GCF_000317375.1 | Microtus ochrogaster |
| GCF_000001635.26 | Mus musculus |
| GCF_900095145.1 | Mus pahari |
| GCF_002201575.1 | Neomonachus schauinslandi |
| GCF_000292845.1 | Ochotona princeps |
| GCF_000260255.1 | Octodon degus |
| GCF_000321225.1 | Odobenus rosmarus divergens |
| GCF_009806435.1 | Oryctolagus cuniculus |
| GCF_000181295.1 | Otolemur garnettii |
| GCF_016772045.2 | Ovis aries |
| GCF_000956105.1 | Propithecus coquereli |
| GCF_003327715.1 | Puma concolor |
| GCF_036323735.1 | Rattus norvegicus |
| GCF_000235385.1 | Saimiri boliviensis boliviensis |
| GCF_000181275.1 | Sorex araneus |
| GCF_000003025.6 | Sus scrofa |
| GCF_000243295.1 | Trichechus manatus latirostris |

### Appendix D. Comparison of Models for *de novo* Gene Annotation

**Table A7.** Comparison of models for *de novo* Gene Annotation

| Model | Architecture (details) | N params, M | Input, Kb | Tokenization | Released |
|---|---|---|---|---|---|
| GENATATOR (GENA large) | Transformer (RMT) + U-Net | 360 | 1000 | BPE | this work |
| GENATATOR (GENA base) | Transformer (RMT) + U-Net | 120 | 32 | BPE | this work |
| GENATATOR (Caduceus PH) | SSM | 15 | 250 | nucleotide | this work |
| GENATATOR (Caduceus PS) | SSM (+RC equivalent) | 15 | 250 | nucleotide | this work |
| Evo2 | SSM | 1000 | 32 | nucleotide | (3) (probing only) |
| AUGUSTUS | HMM | N/A | N/A | 1-bp | (22) |
| Tiberius | CNN + HMM | 8 | 10 | 1-hot | (10) |
| Helixer | CNN + HMM | N/A | 100 | 1-bp | (11) |
| AlphaGenome | CNN + Transformer | 450 | 1000 | 1-bp | (1) |
| SegmentNT | Transformer (RoPE) + U-Net | 500 | 50 | 6-mer | (7) |
| SegmentBorzoi | CNN + U-Net | 323 | 196 | nucleotide | (7) |
| SegmentEnformer | Transformer + U-Net | 379 | 196 | nucleotide | (7) |
| NTv3 (100M) | U-Net + Transformer | 100 | 1000 | nucleotide | (2) |
| NTv3 (650M) | U-Net + Transformer | 650 | 1000 | nucleotide | (2) |

#### Appendix E. Training on embeddings.

**Table A8.** Gene-level metric after training on frozen embeddings of different DNA LM models.

| Model | Chunk length $N_l = 4096$ bp | | | | Chunk length $N_l = 32000$ bp | | | |
|---|---|---|---|---|---|---|---|---|
| | mRNA | | lncRNA | all RNA | mRNA | | lncRNA | all RNA |
| | exon | CDS | exon | exon | exon | CDS | exon | exon |
| GENA base | 4 | 0 | 1 | 5 | 7 | 0 | 2 | 9 |
| Caduceus PH | 0 | 0 | 0 | 0 | 0 | 0 | 0 | 0 |
| Caduceus PS | 0 | 0 | 0 | 0 | 0 | 0 | 0 | 0 |
| Evo2 | 0 | 0 | 0 | 0 | 0 | 0 | 0 | 0 |

#### Appendix F. Clustering of hidden states of the models

*Setup* We extracted final-layer hidden states for ten randomly selected human genes, comprising six mRNA and four lncRNA transcripts. Two model states were analyzed: pretrained HuggingFace (HF) weights and our fine-tuned `GENATATOR` models for both architectures. For `GENA-LM` (BPE tokenization), each token embedding was expanded uniformly across its nucleotide span to obtain one vector per base. Importantly, we intercepted embeddings directly from the RMT backbone prior to the U-Net decoder in order to evaluate the pretrained representation itself. This was necessary because the U-Net component was introduced only in this work and is randomly initialized, as no pretrained version with U-Net exists. Passing embeddings through such a randomly initialized head would risk altering the information contained in the pretrained backbone. For `Caduceus`, weight tying was disabled (`weight_tying=False`) for both HF and fine-tuned states, which doubled the number of trainable parameters (up to 16M parameters). We fit two-dimensional PCA directly to the raw per-base embeddings and then applied $k$-means with $k$=5.

*Homogeneity metric* Let $K$ denote the ground-truth label random variable over exon, intron, CDS, 5′UTR, and 3′UTR, and $C$ the cluster assignment returned by $k$-means. Define

$$H(K) = -\sum_{k} \frac{n_k}{N} \log\left(\frac{n_k}{N}\right), \qquad H(K \mid C) = -\sum_{c}\sum_{k} \frac{n_{c,k}}{N} \log\left(\frac{n_{c,k}}{n_c}\right),$$

where $n_k$ is the count of label $k$, $n_c$ is the size of cluster $c$, $n_{c,k}$ is the number of samples with label $k$ in cluster $c$, and $N$ is the total number of samples. The homogeneity score is

$$\mathrm{h} \;=\; 1 - \frac{H(K \mid C)}{H(K)},$$

with h=1 when $H(K)$=0 (see `sklearn.metrics.homogeneity_score`).

*Selected gene set* The analysis covered the ten human genes listed in Table A9, spanning both coding and non-coding classes and a broad range of transcript lengths.

*Explained variance of PCA* To evaluate how much variance in the embeddings is captured by the leading principal components, we report the explained variance ratios (EVR) of the first two components (Table A10). These values quantify how strongly base identity or higher-order transcript structure dominate the embedding space.

**Table A9.** Gene set used for the embedding analysis. Lengths are transcript lengths in base pairs.

| Gene | Type | Length (bp) |
|---|---|---|
| LOC105375876 | lncRNA | 4,791 |
| CPSF1 | mRNA | 16,281 |
| FDFT1 | mRNA | 36,533 |
| OSER1-DT | lncRNA | 14,964 |
| ERGIC3 | mRNA | 15,580 |
| TPX2 | mRNA | 62,507 |
| NOP56 | mRNA | 5,768 |
| IQANK1 | mRNA | 56,563 |
| LINC02986 | lncRNA | 3,453 |
| LOC107986930 | lncRNA | 140,852 |

**Table A10.** Explained variance ratios (EVR) of the first two PCA components computed directly on per-base embeddings without pooling.

| Model state | $EVR_1$ | $EVR_2$ |
|---|---|---|
| Caduceus PS (HF) | 0.587 | 0.164 |
| Caduceus PS (fine-tuned) | 0.477 | 0.221 |
| GENA LM large (HF) | 0.010 | 0.009 |
| GENA LM large (fine-tuned) | 0.515 | 0.078 |

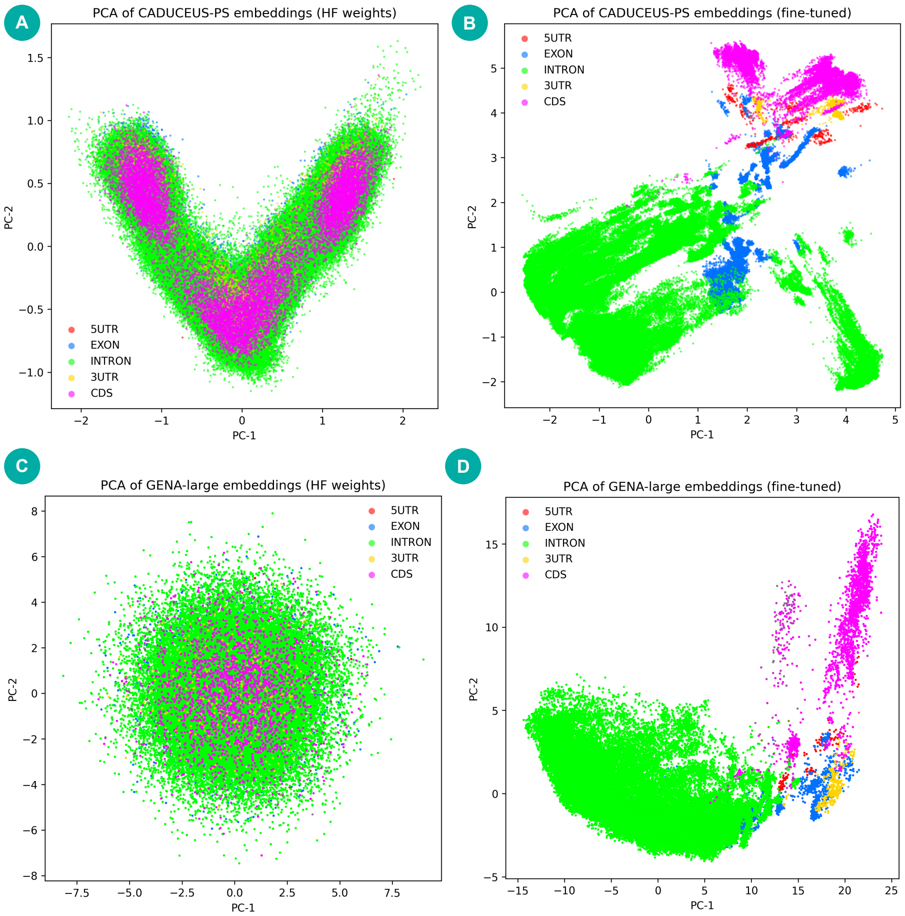


**Fig. A1.** PCA of final-layer embeddings colored by gene-structure labels (5'UTR, EXON, INTRON, 3'UTR, CDS). Panels correspond to `Caduceus PS` with HuggingFace (HF) weights (A), `Caduceus PS` after fine-tuning (B), `GENA LM large` with HF weights (C), and `GENA LM large` after fine-tuning (D).

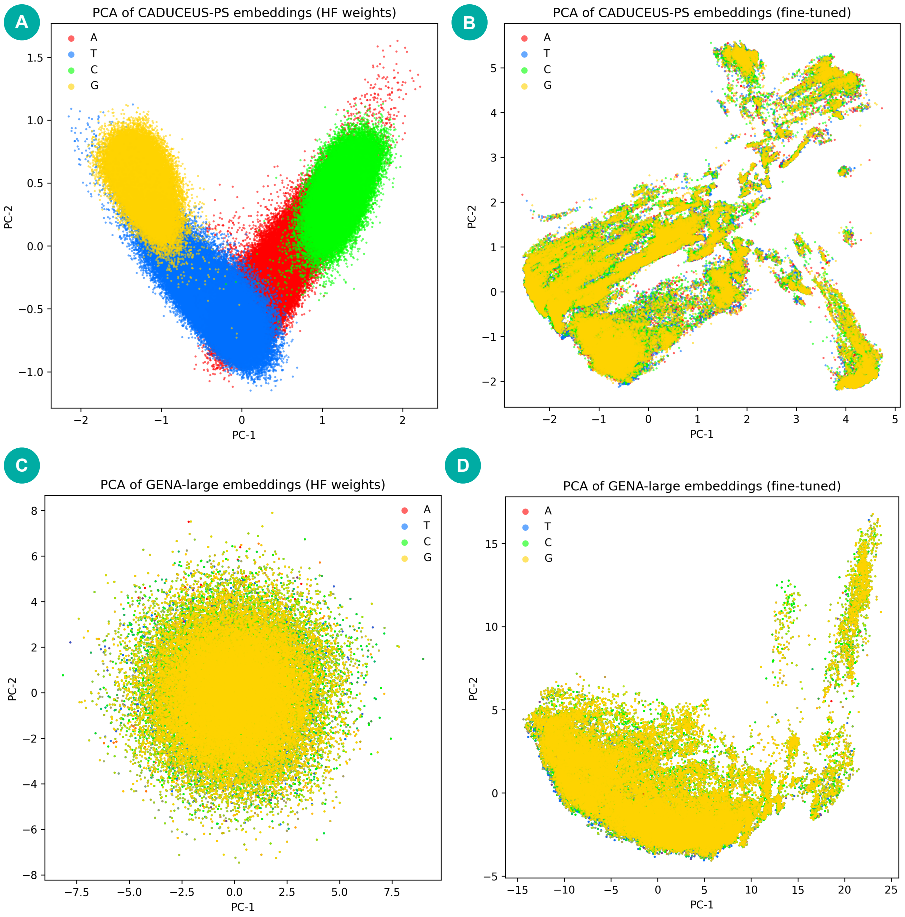


**Fig. A2.** PCA of the same embeddings colored by nucleotide identity (A, T, C, G). Under HF weights, `Caduceus PS` exhibits clear separation by base identity, while fine-tuning reduces base-driven structure and enhances organization by transcript elements.

#### Appendix G. GENATATOR error analysis.

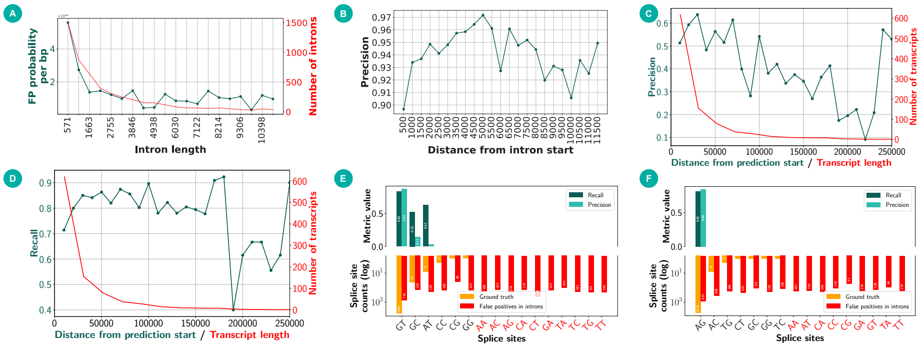


**Fig. A3.** Error analysis provides insights into potential tweaks for improving gene annotation. A-D: Performance metrics as a function of intron length (A), distance from exon-intron boundary (B), and distance from gene sequence start (C-D). B aggregates intron sequences located at a specific distance from the exon-intron boundary. In A and B, the distribution is cropped at the 90th percentile, in C and D at 250Kb. E and F: Precision and recall at predicted intron-exon boundaries, stratified by flanking dinucleotide, separately for left (E) and right (F) intron boundaries, with the distribution of targets shown in red and orange.

#### Appendix H. Computing power requirements.

We intentionally performed vast majority of the experiments on a small dataset using downscaled models (i.e. base `GENA-LM` version instead of large) to save computation time and allow more datasets and architectures to be benchmarked. We believe that providing results of the thorough benchmarking is important background with saves compute for others who is going to develop better models for gene annotation.

Average time and resources required for processing 250 Kbp with the most efficient `GENATATOR` models are provided in the table below. For the whole human chromosome, it takes 15 min using a single A100 GPU and 8.5 GB GRAM. `GENA-based` models can be used even without GPU: with Intel(R) Xeon(R) Platinum 8358 CPU @ 2.60GHz, a single chromosome (chr20, 67Mbp) can be annotated within 3h.

Here, NA indicates that `Caduceus PS` cannot be executed on CPU.

In addition to per-chunk throughput, we also measured end-to-end inference time on a full human chromosome using the same hardware configuration (one NVIDIA A100 80GB). On chromosome 20 of the T2T human genome, the `GENA`-based `GENATATOR` model required approximately 16 minutes to complete

**Table A11.** Runtime and memory usage of different models.

| Model | A100 × 1 Time | A100 × 1 Memory | CPU Time | CPU Memory |
|---|---|---|---|---|
| GENA large | 3.5 s | 8 430 MiB | 42 s | 8 430 MiB |
| Caduceus PS | 1 s | 7 936 MiB | NA | NA |

annotation, while the `Caduceus`-based `GENATATOR` variant completed the same task in about 8 minutes. For comparison, `SegmentNT` (evaluated using its recommended window size of 49,992 bp) required 36 minutes, `Tiberius` completed annotation in 13 minutes, and `AUGUSTUS` required 67 minutes.

#### Appendix I. Models scoring and benchmarking

##### I.1 Processing Predictions

For all models except `Tiberius`, `Helixer` and `AUGUSTUS`, each nucleotide was assigned the class with the highest value from the comparison group. The comparison group is specific to each class: for the exon class, it includes exon and intron. For the CDS class, it includes CDS, intron, 5'UTR, and 3'UTR.

##### I.2 Benchmarking

Predictions were obtained by feeding the model with nucleotide sequences of transcripts. `SegmentNT` is not designed to process very long sequences, so for this model, gene sequences were split into non-overlapping segments of 30 kb and 50 kb. For `NTv3`, input segment lengths of 32 kb and 1 Mb were used. For `SegmentBorzoi`, the input segment length was set to 524,288 nucleotides, and for `SegmentEnformer`, it was set to 196,608 nucleotides, as recommended by the authors.

For `AlphaGenome`, several input sequence lengths are available. Here, we used a segment size of 1 Mb. For the segmentation task, the most suitable track, splice sites, was employed. Exons were defined based on acceptor and donor classes, corresponding to the first and last nucleotide of each exon, respectively. Acceptor-donor pairs were identified in a sliding window from the beginning to the end of the sequence. We evaluated thresholds ranging from 0.1 to 0.9 in increments of 0.1, and for the final results, the best-performing threshold 0.7 was selected.

It is important to note that `SegmentNT` can predict only the exon class, so metrics for the CDS class were obtained by subtracting predictions of 5'UTR and 3'UTR from exon predictions. Finally, `GENATATORs` and `NTv3` are capable of predicting both exons and CDS, so for these models, metrics were calculated across all classes for all genes and transcripts.

##### I.3 PR-AUC

PR-AUC for each class was computed using sequences of all transcripts for each gene, after which the resulting values were averaged.

##### I.4 Interval level metrics

To evaluate the accuracy of exon prediction for each model, sequences of all transcripts for each gene were provided.

##### I.5 CDS-heuristics

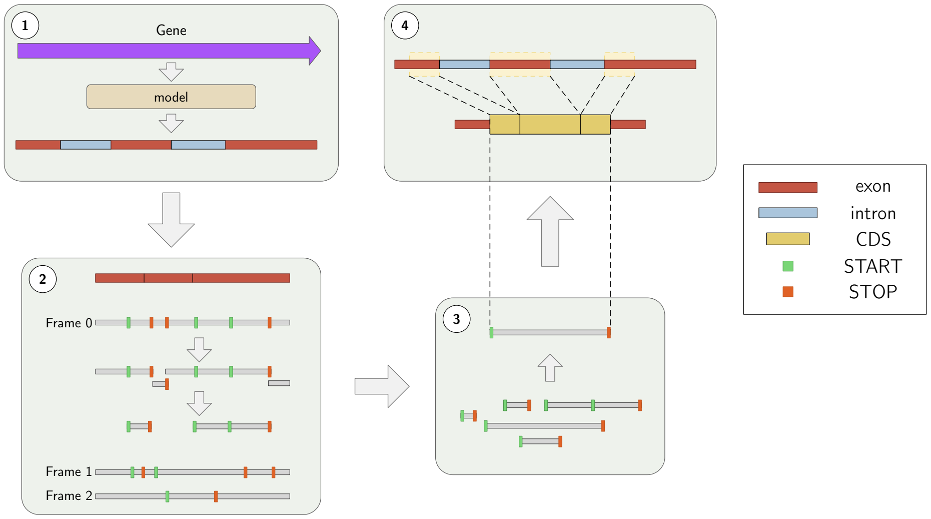


**Fig. A4.** CDS-heuristics workflow. 1. The model predicts the exon-intron structure from the nucleotide sequence. 2. All predicted exons are concatenated, and the resulting nucleotide sequence is translated in all three reading frames. For frame 0, the obtained protein sequence is split at the STOP signal (shown in red). Only fragments containing both a STOP and a START (shown in green) are retained. If necessary, the proteins are clipped so that they begin precisely at the START. The same procedure is applied to the other frames. 3. From the resulting pool of proteins, the longest one is selected. 4. This protein is then mapped back onto the predicted exons to obtain the final CDS.

##### I.6 Gene level

Each model generated predictions based on the each transcript sequences of each gene. Interval-level (exon or CDS) analysis was then performed. If there is a transcript with complete and reciprocal overlap between predicted exons and known exons, the gene was considered to be identified. CDS analysis was performed similarly.

##### I.7 BUSCO

Based on the predictions of each model, the nucleotide sequences of the transcripts were obtained for analysis. After performing the translation operation, the corresponding proteins were obtained and the longest of them was selected. The strand for translation was determined either directly if model outputs it explicitly (`Tiberius` and `AUGUSTUS`), or based on the predicted classes 5'UTR and 3'UTR, using the formula: $(FirstU5 - FirstU3) - (LastU5 - LastU3)$, where $FirstU5$ is the cumulative probability of 5'-UTR class preidction in the first 50 bases, $LastU3$ is the cumulative probability of 3'-UTR class prediction in the last 50 bases, and etc. (for other models). For `AlphaGenome`, the strand corresponding to the gene strand was used. Subsequently, the set of obtained proteins was analyzed using BUSCO.

#### Appendix J. Models scoring and benchmarking.

**Table A12.** Comparison of `GENA base` (Table 1) and `GENA large` in the baseline setup.

| Model | Category | Gene-level |
|---|---|---|
| GENA base | EXON mRNA + lncRNA | 58 |
| | EXON mRNA | 40 |
| | EXON lncRNA | 4 |
| | CDS mRNA | 18 |
| GENA large | EXON mRNA + lncRNA | 74 |
| | EXON mRNA | 54 |
| | EXON lncRNA | 20 |
| | CDS mRNA | 5 |

**Table A13.** PR-AUC benchmark, related to Figure 2.

| Model | Caduceus PS | GENA large RMT | NTv3 (100M) | NTv3 (650M) | SegmentNT | SegmentNT multi_species | SegmentEnformer | SegmentBorzoi |
|---|---|---|---|---|---|---|---|---|
| Input | 250k | 1 Mb | 1 Mb | 1 Mb | 50 kb | 50 kb | 196 kb | 524 kb |
| Mean | **0.8315** | 0.7815 | 0.7157 | 0.7225 | 0.7131 | 0.7070 | 0.5329 | 0.5200 |
| 5UTR | **0.7086** | 0.6094 | 0.4783 | 0.4912 | 0.4648 | 0.4502 | 0.1910 | 0.1914 |
| Exon | **0.9652** | 0.9153 | 0.8174 | 0.8129 | 0.8231 | 0.8237 | 0.6954 | 0.6755 |
| Intron | **0.9705** | 0.9626 | 0.9359 | 0.9436 | 0.9265 | 0.9232 | 0.8391 | 0.8382 |
| 3UTR | **0.7675** | 0.7116 | 0.6424 | 0.6546 | 0.6378 | 0.6310 | 0.4060 | 0.3749 |
| CDS | **0.7457** | 0.7087 | 0.7045 | 0.7104 | - | - | - | - |

**Table A14.** BUSCO mammalia completeness computed on hold-out gene set (human chromosome 20). Related to A14.

| Model | Input | Complete | Fragmented |
|---|---|---|---|
| GENA large RMT | 1 Mb | 224 | 27 |
| Caduceus PS | 250 kb | 222 | 30 |
| Tiberius | | **247** | 6 |
| AUGUSTUS | | 196 | 30 |
| SegmentNT | 30 kb | 174 | 46 |
| SegmentNT | 50 kb | 176 | 45 |
| SegmentNT_multi_species | 30 kb | 170 | 45 |
| SegmentNT_multi_species | 50 kb | 170 | 48 |
| SegmentBorzoi | 524 kb | 36 | 33 |
| SegmentEnformer | 196 kb | 31 | 31 |
| NT_v3 (100M) | 32 kb | 0 | 0 |
| NT_v3 (100M) | 1 Mb | 0 | 0 |
| NT_v3 (650M) | 32 kb | 0 | 0 |
| NT_v3 (650M) | 1 Mb | 0 | 0 |
| Helixer | | 135 | 4 |
| AlphaGenome | 1 Mb | 212 | 30 |
| Ground True | | 275 | 3 |

**Table A15.** Interval-level benchmark. Related to Figure 2.

| Model | Input | Gene type | Class | Precision | Recall | F1 |
|---|---|---|---|---|---|---|
| GENA large RMT | 1 Mb | mRNA | EXON | 0.85 | 0.85 | 0.85 |
| | | mRNA | CDS | 0.83 | 0.67 | 0.74 |
| | | lncRNA | EXON | 0.53 | 0.51 | 0.52 |
| | | All RNA | EXON | 0.82 | 0.82 | 0.82 |
| Caduceus PS | 250 kb | mRNA | EXON | **0.89** | **0.87** | **0.88** |
| | | mRNA | CDS | **0.85** | **0.73** | **0.79** |
| | | lncRNA | EXON | **0.62** | **0.52** | **0.56** |
| | | All RNA | EXON | **0.87** | **0.84** | **0.85** |
| Tiberius | 250 kb | mRNA | EXON | 0.74 | 0.49 | 0.59 |
| | | mRNA | CDS | **0.85** | 0.63 | 0.73 |
| | | lncRNA | EXON | 0.22 | 0.01 | 0.02 |
| | | All RNA | EXON | 0.73 | 0.45 | 0.56 |
| AUGUSTUS | 250 kb | mRNA | EXON | 0.62 | 0.59 | 0.61 |
| | | mRNA | CDS | 0.66 | 0.67 | 0.66 |
| | | lncRNA | EXON | 0.08 | 0.02 | 0.03 |
| | | All RNA | EXON | 0.61 | 0.54 | 0.57 |
| Helixer | 250 kb | mRNA | EXON | 0.62 | 0.56 | 0.59 |
| | | mRNA | CDS | 0.66 | 0.62 | 0.64 |
| | | lncRNA | EXON | 0.19 | 0.02 | 0.03 |
| | | All RNA | EXON | 0.62 | 0.52 | 0.56 |
| AlphaGenome | 1 Mb | mRNA | EXON | 0.85 | 0.79 | 0.82 |
| | | mRNA | CDS | 0.70 | 0.72 | 0.71 |
| | | lncRNA | EXON | 0.34 | 0.15 | 0.20 |
| | | All RNA | EXON | 0.83 | 0.73 | 0.78 |
| SegmentNT | 30 kb | mRNA | EXON | 0.46 | 0.74 | 0.57 |
| | | mRNA | CDS | 0.38 | 0.69 | 0.49 |
| | | lncRNA | EXON | 0.08 | 0.10 | 0.09 |
| | | All RNA | EXON | 0.43 | 0.69 | 0.53 |
| SegmentNT | 50 kb | mRNA | EXON | 0.48 | 0.74 | 0.58 |
| | | mRNA | CDS | 0.40 | 0.68 | 0.50 |
| | | lncRNA | EXON | 0.09 | 0.10 | 0.09 |
| | | All RNA | EXON | 0.46 | 0.68 | 0.55 |
| SegmentNT_multi_species | 30 kb | mRNA | EXON | 0.41 | 0.74 | 0.53 |
| | | mRNA | CDS | 0.34 | 0.68 | 0.46 |
| | | lncRNA | EXON | 0.10 | 0.12 | 0.11 |
| | | All RNA | EXON | 0.39 | 0.68 | 0.50 |
| SegmentNT_multi_species | 50 kb | mRNA | EXON | 0.42 | 0.73 | 0.53 |
| | | mRNA | CDS | 0.35 | 0.67 | 0.46 |
| | | lncRNA | EXON | 0.12 | 0.12 | 0.12 |
| | | All RNA | EXON | 0.40 | 0.68 | 0.51 |
| NT_v3 (100M) | 32 kb | mRNA | EXON | 0.09 | 0.23 | 0.13 |
| | | mRNA | CDS | 0.07 | 0.11 | 0.09 |
| | | lncRNA | EXON | 0.01 | 0.05 | 0.02 |
| | | All RNA | EXON | 0.08 | 0.21 | 0.12 |
| NT_v3 (100M) | 1 Mb | mRNA | EXON | 0.33 | 0.67 | 0.44 |
| | | mRNA | CDS | 0.20 | 0.32 | 0.25 |
| | | lncRNA | EXON | 0.02 | 0.07 | 0.03 |
| | | All RNA | EXON | 0.29 | 0.62 | 0.39 |
| NT_v3 (650M) | 32 kb | mRNA | EXON | 0.16 | 0.24 | 0.19 |
| | | mRNA | CDS | 0.07 | 0.09 | 0.08 |
| | | lncRNA | EXON | 0.03 | 0.06 | 0.04 |
| | | All RNA | EXON | 0.14 | 0.23 | 0.18 |
| NT_v3 (650M) | 1 Mb | mRNA | EXON | 0.47 | 0.72 | 0.56 |
| | | mRNA | CDS | 0.20 | 0.29 | 0.24 |
| | | lncRNA | EXON | 0.09 | 0.14 | 0.11 |
| | | All RNA | EXON | 0.43 | 0.67 | 0.52 |

**Table A16.** Gene level metrics computed on a gene set assembled from 14 animal species. Metrics are calculated for protein-coding and non-coding genes in a gene set from a single chromosome for each species. MRCA MYA - million years from most recent common ancestor with Homo sapiens.

| Species | MRCA (MYA) | Chromosome | Gene type | Class | Caduceus PS | GENA large RMT | Tiberius | AUGUSTUS | Ground true |
|---|---|---|---|---|---|---|---|---|---|
| Arabidopsis thaliana | 1530 | NC_003075.7 | mRNA | EXON | **1355** | 1330 | 0 | 298 | 4180 |
| | | | | CDS | 1462 | 1476 | 594 | **2286** | |
| | | | lncRNA | EXON | **310** | 307 | 0 | 22 | 513 |
| | | | all RNA | EXON | **1665** | 1637 | 0 | 320 | 4693 |
| Saccharomyces cerevisiae S288C | 1275 | NC_001136.10 | mRNA | EXON | **733** | 697 | 0 | 0 | 766 |
| | | | | CDS | **725** | 689 | 0 | 358 | |
| | | | lncRNA | EXON | | | | | 0 |
| | | | all RNA | EXON | **733** | 697 | 0 | 0 | 766 |
| Anopheles funestus | 686 | NC_064599.1 | mRNA | EXON | **1765** | 1606 | 0 | 38 | 4821 |
| | | | | CDS | **2257** | 206 | 993 | 1921 | |
| | | | lncRNA | EXON | 23 | **27** | 0 | 0 | 243 |
| | | | all RNA | EXON | **1788** | 1633 | 0 | 38 | 5064 |
| Drosophila melanogaster | 686 | NT_033779.5 | mRNA | EXON | **1291** | 1201 | 0 | 64 | 2657 |
| | | | | CDS | 1387 | 1323 | 866 | **1462** | |
| | | | lncRNA | EXON | **280** | 261 | 0 | 0 | 526 |
| | | | all RNA | EXON | **1571** | 1462 | 0 | 64 | 3183 |
| Danio rerio | 429 | NC_133178.1 | mRNA | EXON | **574** | 467 | 0 | 0 | 1345 |
| | | | | CDS | **614** | 537 | 453 | 352 | |
| | | | lncRNA | EXON | 83 | **101** | 0 | 0 | 344 |
| | | | all RNA | EXON | **657** | 568 | 0 | 0 | 1689 |
| Mugil cephalus | 429 | NC_061770.1 | mRNA | EXON | **834** | 737 | 0 | 0 | 2119 |
| | | | | CDS | **900** | 817 | 781 | 570 | |
| | | | lncRNA | EXON | **108** | 94 | 0 | 0 | 293 |
| | | | all RNA | EXON | **942** | 831 | 0 | 0 | 2412 |
| Paralichthys olivaceus | 429 | NC_091093.1 | mRNA | EXON | **399** | 345 | 0 | 0 | 944 |
| | | | | CDS | **438** | 396 | 409 | 278 | |
| | | | lncRNA | EXON | **27** | 25 | 0 | 0 | 129 |
| | | | all RNA | EXON | **426** | 370 | 0 | 0 | 1073 |
| Xenopus laevis | 352 | NC_054386.1 | mRNA | EXON | **615** | 484 | 0 | 4 | 1463 |
| | | | | CDS | **694** | 545 | 484 | 164 | |
| | | | lncRNA | EXON | **60** | 48 | 0 | 0 | 161 |
| | | | all RNA | EXON | **675** | 532 | 0 | 4 | 1624 |
| Anas platyrhynchos | 319 | NC_092591.1 | mRNA | EXON | **586** | 491 | 0 | 0 | 1002 |
| | | | | CDS | **628** | 543 | 617 | 280 | |
| | | | lncRNA | EXON | **67** | 48 | 0 | 0 | 412 |
| | | | all RNA | EXON | **653** | 539 | 0 | 0 | 1414 |
| Gallus gallus | 319 | NC_052536.1 | mRNA | EXON | **574** | 485 | 0 | 0 | 1036 |
| | | | | CDS | 614 | 541 | **616** | 300 | |
| | | | lncRNA | EXON | **59** | 40 | 0 | 0 | 314 |
| | | | all RNA | EXON | **633** | 525 | 0 | 0 | 1350 |
| Taeniopygia guttata | 319 | NC_133030.1 | mRNA | EXON | **528** | 467 | 0 | 0 | 976 |
| | | | | CDS | 571 | 514 | **594** | 262 | |
| | | | lncRNA | EXON | **62** | 43 | 0 | 0 | 245 |
| | | | all RNA | EXON | **590** | 510 | 0 | 0 | 1221 |
| Bubalus bubalis | 94 | NC_059174.1 | mRNA | EXON | **774** | 757 | 0 | 12 | 1239 |
| | | | | CDS | **843** | 839 | 753 | 240 | |
| | | | lncRNA | EXON | **87** | 70 | 0 | 0 | 331 |
| | | | all RNA | EXON | **861** | 827 | 0 | 12 | 1570 |
| Panthera tigris | 94 | NC_056673.1 | mRNA | EXON | **695** | 694 | 0 | 30 | 1136 |
| | | | | CDS | 734 | **747** | 670 | 139 | |
| | | | lncRNA | EXON | **86** | 68 | 0 | 0 | 284 |
| | | | all RNA | EXON | **781** | 762 | 0 | 30 | 1420 |
| Tursiops truncatus | 94 | NC_047043.1 | mRNA | EXON | **591** | 523 | 0 | 18 | 1079 |
| | | | | CDS | **635** | 597 | 603 | 199 | |
| | | | lncRNA | EXON | **52** | 31 | 0 | 0 | 214 |
| | | | all RNA | EXON | **643** | 554 | 0 | 18 | 1293 |
| Pan troglodytes | 6.4 | NC_072417.2 | mRNA | EXON | 780 | **782** | 0 | 13 | 1304 |
| | | | | CDS | 832 | **859** | 786 | 228 | |
| | | | lncRNA | EXON | **74** | 59 | 0 | 0 | 284 |
| | | | all RNA | EXON | **854** | 841 | 0 | 13 | 1588 |
| Homo sapiens | 0 | NC_060944.1 | mRNA | EXON | **377** | 347 | 0 | 6 | 546 |
| | | | | CDS | **392** | 362 | 379 | 111 | |
| | | | lncRNA | EXON | **111** | 95 | 0 | 0 | 434 |
| | | | all RNA | EXON | **488** | 442 | 0 | 6 | 980 |

#### Appendix K. Homology Exclusion Experiment in *S. cerevisiae.*

To ensure that the performance of GENATATORs in yeast is not attributable to residual homology with mammalian training data, we performed a stringent control. All 766 annotated protein-coding genes from S. cerevisiae chromosome NC_001136.10 were compared to the full proteomes of the 39 mammalian species used during training (1,827,441 proteins in total) using BLASTP (E-value cutoff 1e–05). Every yeast gene with at least one significant hit was excluded, resulting in a filtered set of 270 genes without detectable protein-level similarity to the training data.

We then evaluated gene-level reconstruction accuracy on this filtered set. Results are summarized in Table A17.

**Table A17.** Gene-level reconstruction on *S. cerevisiae* genes without detectable protein-level homology to mammals.

| Model | Gene level |
|---|---|
| Caduceus PS | 263 |
| GENA large | 250 |
| AUGUSTUS | 113 |
| Ground truth | 270 |

Even under these stringent conditions, GENATATORs recovered over 250 genes - more than twice the number recovered by AUGUSTUS, which was run with a species-specific HMM profile for *S. cerevisiae.* These findings demonstrate that the observed performance cannot be explained by homology leakage, but instead reflects the models' ability to capture general splice and coding sequence patterns transferable across kingdoms.

#### Appendix L. Declaration of LLM usage.

Large Language Models (LLMs) were used solely to improve the readability and clarity of the manuscript text. No parts of the analysis, results, or conclusions were generated by LLMs.
